# Mangroves are an overlooked hotspot of insect diversity despite low plant diversity

**DOI:** 10.1101/2020.12.17.423191

**Authors:** Darren Yeo, Amrita Srivathsan, Jayanthi Puniamoorthy, Foo Maosheng, Patrick Grootaert, Lena Chan, Benoit Guénard, Claas Damken, Rodzay A. Wahab, Ang Yuchen, Rudolf Meier

## Abstract

We here compare the tropical arthropod fauna across a freshwater swamp and six different forest types (rain-, swamp, dry-coastal, urban, freshwater swamp, mangroves) based on 140,000 barcoded specimens belonging to *ca*. 8,500 species. Surprisingly, we find that mangroves, a globally imperiled habitat that had been expected to be species-poor for insects, are an overlooked hotspot for insect diversity despite having low plant diversity. Our study reveals a species-rich mangrove insect fauna (>3,000 species) that is distinct (>50% of species are mangrove-specific), and has high species turnover across Southeast and East Asia. For most habitats, plant diversity is a good predictor for insect diversity, but mangroves are an exception and compensate for a comparatively low number of phytophagous and fungivorous insect species by supporting an unusually rich community of predators whose larvae feed in the productive mudflats. For the remaining tropical habitats, the insect communities have diversity patterns that are largely congruent across guilds. The discovery of such a sizeable and distinct insect fauna in a globally threatened habitat underlines how little is known about global insect biodiversity.

## Introduction

Insects are currently experiencing anthropogenic biodiversity meltdowns with declines having attracted much attention [1–4] and controversy [5–10]. The controversy is largely due to the paucity of high-quality, quantitative data with sufficient taxonomic resolution for arthropods. The same paucity is also responsible for imprecise estimates of global animal species richness [11, 12] and the poor understanding of geographic and temporal species turnovers [13–15]. These knowledge gaps are likely to threaten the health of whole ecosystems given that arthropods provide important ecosystem services [3, 16–19], contribute much of the terrestrial animal biomass (~10%) [20] and are yet frequently ignored in habitat assessments. The lack of baseline data is particularly worrisome at a time when tropical ecosystems are heavily impacted by habitat conversion and global change [21].

The situation is particularly dire for the species-rich tropics, for which few comprehensive surveys have been conducted [22–24]. For example, only three of 73 studies in a recent review of insect declines involved tropical sites [8]. Furthermore, tropical insect surveys have traditionally focused on rainforests [24], with other habitats being largely neglected. Mangrove forests are a prime example of a tropical habitat for which the insect fauna is poorly characterized. Mangroves used to cover more than 200,000 km^2^ of the global coastline [25], but have been experiencing an annual area loss of 1-2% [25, 26]. Indeed, the losses of mangroves far exceed those of more high-profile ecosystems such as rainforests and coral reefs [26]. Unfortunately, these losses are further exacerbated by climate change [27], with some simulations predicting a further reduction by 46–59% for all global coastal wetlands by the year 2100 [28]. This is particularly worrying as mangrove ecosystems sequestrate a particularly large amoung of carbon per hectare [29]. These changes will not only endanger entire ecosystems that provide essential ecosystem services [30–32], but also threaten the survival of numerous mangrove species with unique adaptations. Mangrove specialists with such adaptations are well known for vertebrates and vascular plants [33, 34], but the invertebrate diversity is poorly known.

One reason why the mangrove insect fauna may have have received little attention is low plant diversity. Tropical arthropod diversity is usually positively correlated with plant diversity [23, 24, 35] which suggested that mangroves would not be species-rich for insects and provide few insights into one of the key questions in insect biodiversity, *viz*. understanding whether insect herbivores drive high plant diversity in the tropics [36–38] or *vice versa* [22, 39]. Arguably, the traditional focus on this question may have had the undesirable side-effect that the insect fauna of habitats with low plant diversity received little interest. Yet, many of these habitats are threatened with destruction. The few existing studies of mangrove insects focused on specific taxa [40–42], only identified specimens to higher taxonomic levels [43–45], and/or lacked quantitative comparison with the insect fauna of adjacent habitats. Given these shortcomings, it may not surprise that these studies yielded conflicting results [44, 46, 47] with some arguing that high salinity and/or low plant diversity [33, 44, 46] were responsible for a comparatively poor insect fauna, while others reporting high levels of species diversity and specialization [47].

Here, we present the results of a comprehensive study of species richness and turnover of arthropods across multiple tropical habitats. The assessment is based on >140,000 barcoded specimens obtained over >4 years from mangroves, rainforests, swamp forests, disturbed secondary urban forests, dry coastal forests, and freshwater swamps in Singapore (Fig. S1). In addition, we assess the species richness and turnover of mangrove insects across East and Southeast Asia by including samples from Brunei, Thailand, and Hong Kong. Specifically, our study (1) estimates mangrove insect diversity, (2) evaluates the distinctness in reference to five different forest habitats, (3) analyzes the biodiversity patterns by ecological guild, and (4) determines species turnover across larger geographic scales. Most of the work was carried out in Singapore because it has a large variety of different habitats that occur within 40km on a small island (724 km^2^) that lacks major physical barriers. In addition, all habitats have experienced similar levels of habitat degradation or loss (>95% overall loss of original vegetation cover [48]; *ca*. 90% loss of rainforest [49]; *ca*. 93% loss of swamp forest [50]; 91% loss for mangroves [51]).

A thorough assessment of insect biodiversity requires dense sampling over an extended period of time [52–54]. We sampled 107 sites in Singapore using Malaise traps. All samples were subsequently sorted to 13 arthropod orders (Table S2) with Diptera and Hymenoptera contributing the largest number of specimens (>75%: Table S2, see Fig. S2 for species composition). Given the large number of specimens collected, we only barcoded subsets that represented different taxa and ecological guild (see Materials and Methods; Table S3). The individually NGS-barcoded 140,000 specimens [55] were grouped into putative species, which then allowed for estimating species richness and abundance [56–58]. Contrary to expectations, we demonstrate that mangrove forests have a very distinct and rich insect fauna. In addition, the species turnover for all habitats in Singapore and the different mangrove sites in Asia is high.

## Results

### Species delimitation based on NGS barcodes

We obtained 143,807 313-bp *cox1* barcodes in total, which clustered into 8,256–8,903 molecular operationally taxonomic units (mOTUs, henceforth referred to as species) using objective clustering [59] at different p-distance thresholds that allow for assessing the sensitivity of results to clustering thresholds (2–4%; Table S6). A further assessment of robustness was based on using an alternative species delimitation algorithm, USEARCH [60], which yielded similar species richness estimates of 8,520–9,315 species using the identity (--id) parameters 0.96–0.98. Most species boundaries were stable, with species numbers only varying by <12% across species delimitation techniques and parameters. We hence used the species generated via objective clustering at 3% p-distance for the analyses (but see supplementary data Fig. S3 for results obtained with 2% and 4%). We initially analysed the core dataset consisting of 62,066 Diptera and Hymenoptera specimens (4,002 species at 3% p-distance; see Table 2). Thereafter, we broadened the analysis to the full dataset (143,807, see Materials & Methods for sampling details) including data for all orders and sites. We found that the conclusions for mangrove insect diversity were consistent between the datasets. Therefore, subsequent analyses were based on the full dataset, but the corresponding results for the core dataset are presented in Figures S4 and S5.

### Alpha-diversity across habitats

We initially rarefied the species richness curves by sample coverage [61] (Fig. 1) for each habitat, as well as by the number of specimens processed (Fig. S3). Alpha-diversity comparisons were made at the rarefaction point with the lowest coverage/number of specimens (i.e., swamp forest in Fig. 1, *top*). The initial analysis treated mangroves as a single habitat type. The species diversity of mangroves (1,102.5 ± 10.8 species) was ca. 50-60% (core dataset: 40-50%) of the rarefied species richness of adjacent tropical primary/secondary forest (2,188.4 ± 42.6 species) and swamp forest sites (1,809 species) (Fig. 1a). A separate analysis of two major mangrove sites (PU & SB) revealed that they have similar species richness as the freshwater swamp site after rarefaction (Fig. 1b). The species richness of a third mangrove site (SMO) was lower and more similar to the richness of an urban forest site. A newly regenerated mangrove (SMN), adjacent to an old-growth mangrove (SMO) had much lower species richness.

**Figure 1.**
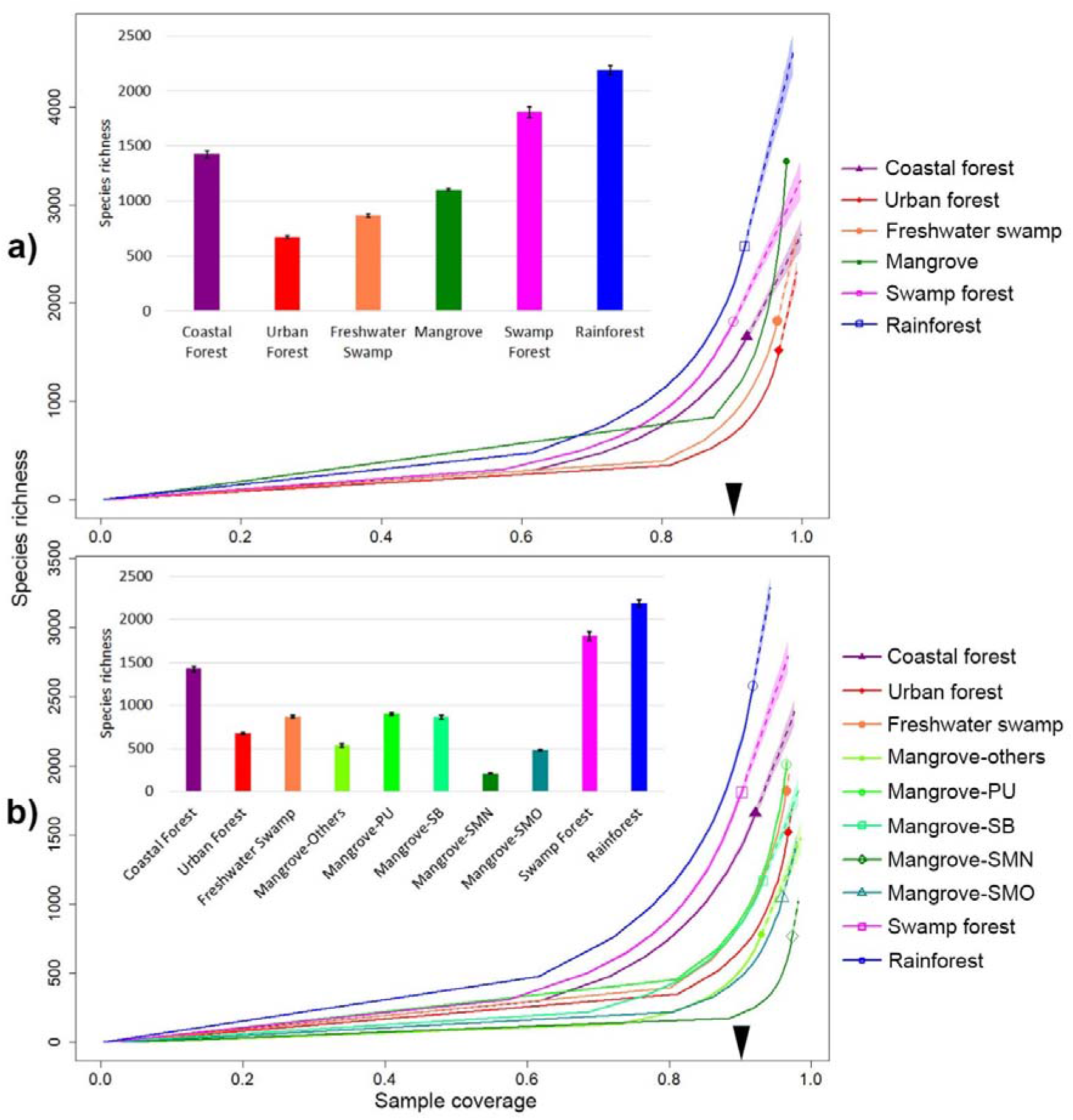
Insect alpha-diversity across tropical forest habitats. (a) Mangroves treated as one habitat; (b) Comparison of mangrove sites: Pulau Ubin (PU), Sungei Buloh (SB), Pulau Semakau old-growth (SMO), Pulau Semakau new-growth (SMN); solid lines = rarefaction; dotted = extrapolations. The arrow on the x-axis indicates the point of rarefaction where species richness comparisons were made, which is reflected in the bar charts with associated 95% confidence intervals.

We also attempted to estimate species richness using extrapolation via frequency ratio [62] and combined non-parametric estimators (CNE) [63], with the former being widely used to estimate microbial diversity [64, 65]. Both estimators suggest that mangroves are even more species-rich than rainforest and swamp forest habitats (Fig. S4) with coastal- and urban forests yielding even higher estimates. However, the confidence intervals are very large and the observed species richness is only 21 – 90% of the estimated richness, which indicates that our data remain insufficient for comparing species richness based on extrapolation [66].

### Species turnover across habitats

Arthropod communities from most habitats are well separated on NMDS plots. This includes the mangrove communities. (Fig. 2) even though several mangrove sites (PU, SB, SM) are geographically further from each other (>30 km) than from the other habitat types (Fig. S1). This separation by habitat is also evident in the core dataset (Fig. S6). These patterns are also observed when the data are split into three taxon sets: (1) Diptera, (2) Hymenoptera, (3) remaining arthropod taxa (Fig. 2b). This habitat-driven structure was verified with a multivariate analysis of variance (MANOVA) test on the NMDS1 & NMDS2 coordinates, with habitat being a highly significant variable (p < 0.001) in explaining the variance in the combined NMDS coordinate distributions and each coordinate separately. These results are also robust to the removal of rare species (Fig. 2a; applies to NMDS plots and MANOVA). Only 48 (0.6%) of the 8,572 putative species in the species turnover analysis are found in all habitat types while 5,989 (69.9%) are only in a single type (Table S7); within the mangroves, 50.2% of the 3,557 species are only known from the mangrove habitat. The habitat type the mangroves share the most species with is the coastal forest (873 of 3,557 species, 24.5%). When rare species are removed (<10 specimens), 481 of the remaining 1,773 species (27.1%) are found in a single habitat while only 48 (2.7%) are found in all (Table S6); i.e., even after excluding rare species, a large proportion of the insect communities are putative habitat specialists.

**Figure 2.**
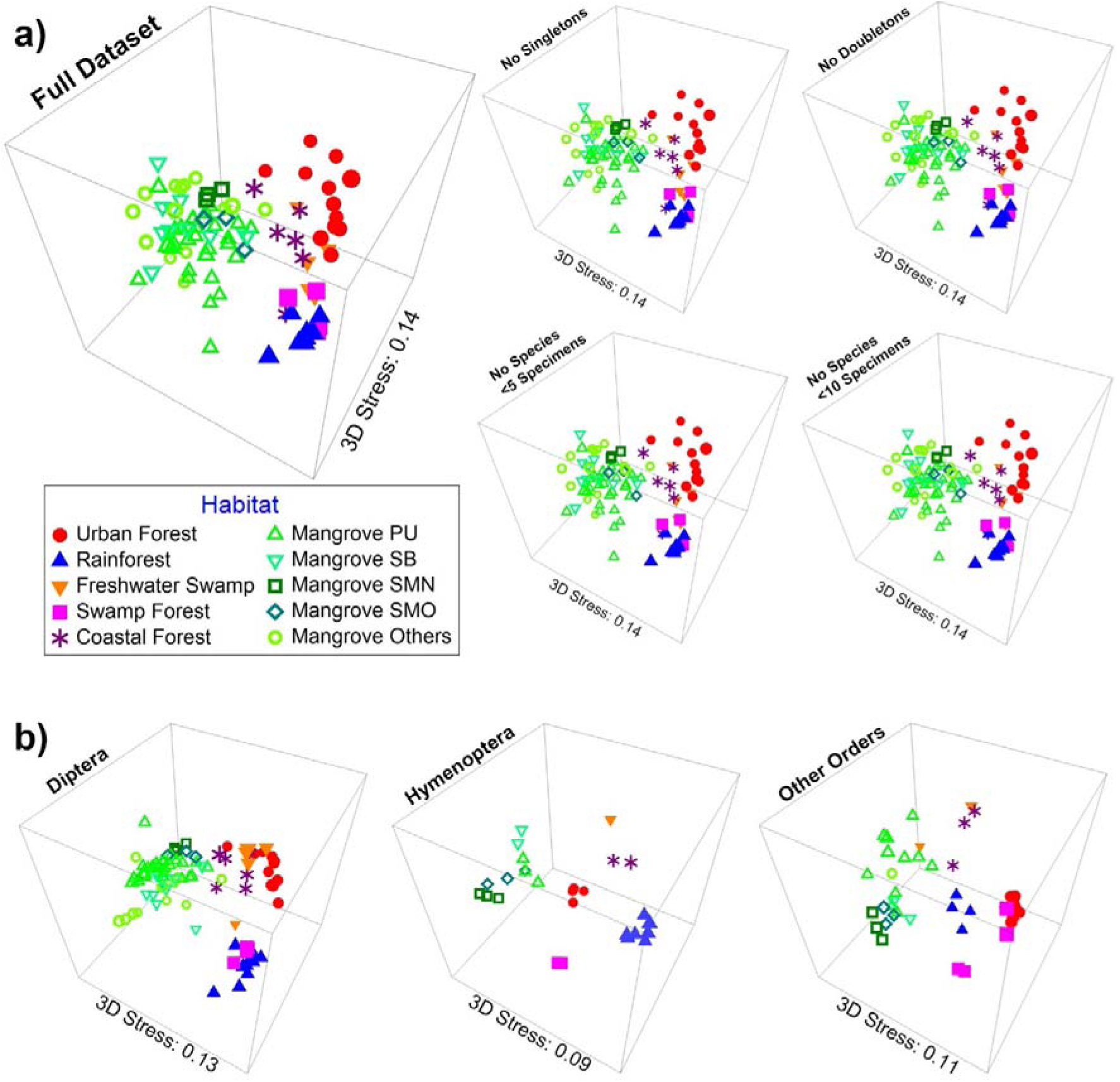
Insect communities across tropical forest habitats are distinct based on Bray-Curtis distances illustrated on 3D NMDS plots, regardless of whether (a) rare species are removed or (b) the data are split into different taxonomic groups.

The dissimilarity of the habitat-specific communities was confirmed via tests with *mvabund* [67], which finds that the variable “habitat” was highly significant (p < 0.001) in determining community structure, even when rare species were excluded. This differentiation is further supported by ANOSIM tests (Table 1A), which find significant differences between communities in both global (P = 0.001, R = 0.784) and pairwise habitat comparisons (P = 0.001 – 0.019, R = 0.341 – 0.983). The only exception are the coastal and urban forests (P = 0.079, R = 0.172) which may be due to the close proximity of Pulau Ubin coastal forest sites to urban settlements (Fig. S1). Note that a SIMPER analysis (Table 1B) finds a substantial number of shared species between the rainforest and swamp forest sites (13.88%). Both sites are in close geographic proximity (<5km; Fig. S1) and the within-habitat values for both sites are fairly high (rainforest = 29.59%, swamp forest = 31.10%). ANOSIM and SIMPER results are again robust to the removal of rare species (Tables S8 & S9) and the ANOSIM p-values for most comparisons are significant even according to re-defined statistical criteria for unexpected or new results (p < 0.005) [68]. The observed dissimilarity was largely due to species turnover with the turnover component (0.898) greatly outweighing nestedness (0.048; Table 1C & S10). This was similarly observed in most pairwise comparisons of habitats (turnover = 0.704 – 0.956, nestedness = 0.001 – 0.102). The only exception was mangroves and coastal forests (turnover = 0.658, nestedness = 0.254) which are in close geographic proximity on Pulau Ubin (Fig. 1).

**Table 1.**
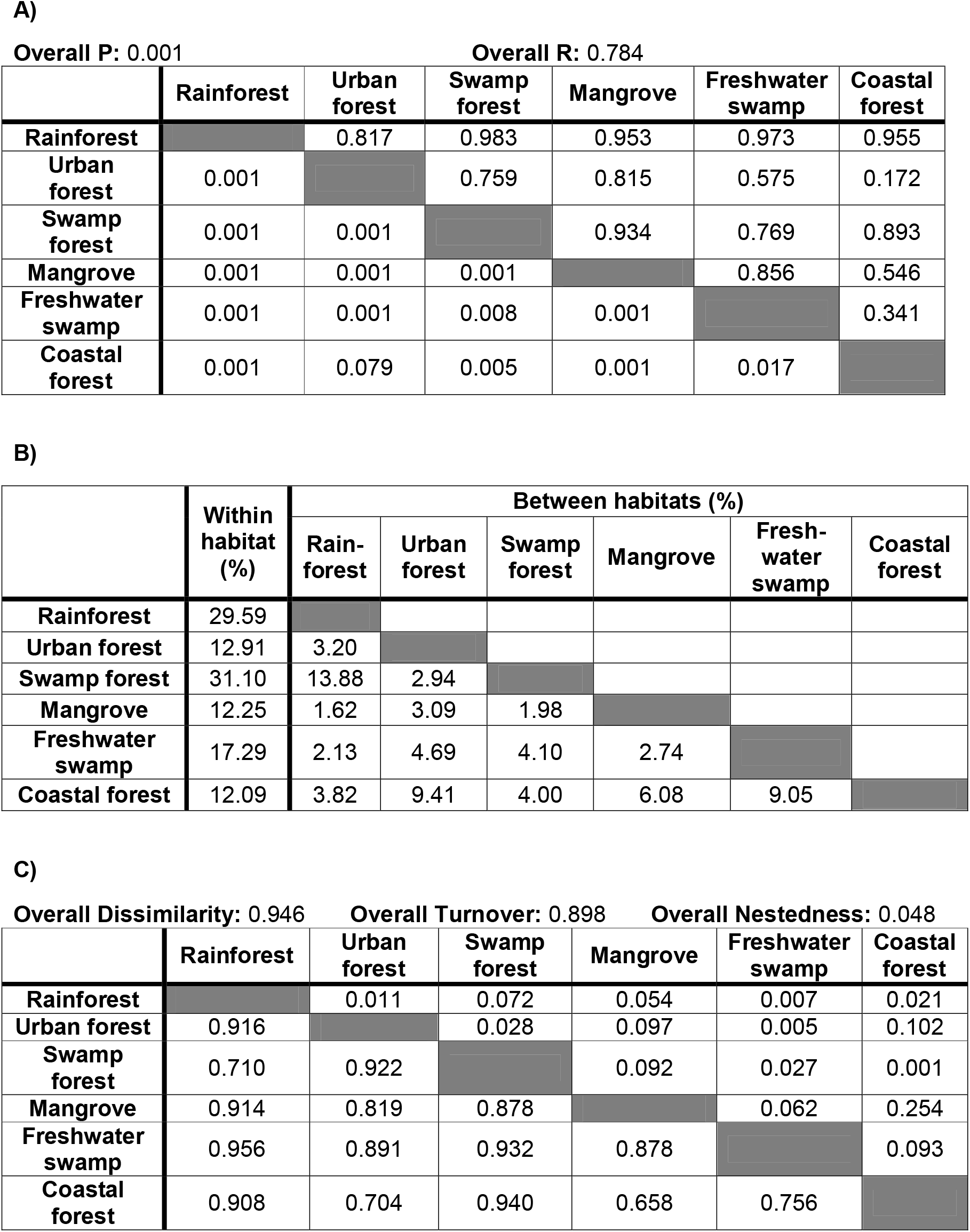
Species turnover across habitats. (A) Distinctness of communities in each habitat type as assessed with ANOSIM (pairwise p-value below and R-statistics above diagonal. (B) Distinctness of communities in each habitat type as assessed with SIMPER. (C) Species turnover and nestedness analysis (pairwise turnover values below and nestedness above diagonal).

**Overall R: 0.784**
|  | Rainforest | Urban forest | Swamp forest | Mangrove | Freshwater swamp | Coastal forest |
| --- | --- | --- | --- | --- | --- | --- |
| Rainforest |  | 0.817 | 0.983 | 0.953 | 0.973 | 0.955 |
| Urban forest | 0.001 |  | 0.759 | 0.815 | 0.575 | 0.172 |
| Swamp forest | 0.001 | 0.001 |  | 0.934 | 0.769 | 0.893 |
| Mangrove | 0.001 | 0.001 | 0.001 |  | 0.856 | 0.546 |
| Freshwater swamp | 0.001 | 0.001 | 0.008 | 0.001 |  | 0.341 |
| Coastal forest | 0.001 | 0.079 | 0.005 | 0.001 | 0.017 |  |

**B)**
|  | Within habitat (%) | Between habitats (%) |  |  |  |  |  |
| --- | --- | --- | --- | --- | --- | --- | --- |
|  |  | Rain-forest | Urban forest | Swamp forest | Mangrove | Fresh-water swamp | Coastal forest |
| Rainforest | 29.59 |  |  |  |  |  |  |
| Urban forest | 12.91 | 3.20 |  |  |  |  |  |
| Swamp forest | 31.10 | 13.88 | 2.94 |  |  |  |  |
| Mangrove | 12.25 | 1.62 | 3.09 | 1.98 |  |  |  |
| Freshwater swamp | 17.29 | 2.13 | 4.69 | 4.10 | 2.74 |  |  |
| Coastal forest | 12.09 | 3.82 | 9.41 | 4.00 | 6.08 | 9.05 |  |

**Overall Nestedness: 0.048**
|  | Rainforest | Urban forest | Swamp forest | Mangrove | Freshwater swamp | Coastal forest |
| --- | --- | --- | --- | --- | --- | --- |
| Rainforest |  | 0.011 | 0.072 | 0.054 | 0.007 | 0.021 |
| Urban forest | 0.916 |  | 0.028 | 0.097 | 0.005 | 0.102 |
| Swamp forest | 0.710 | 0.922 |  | 0.092 | 0.027 | 0.001 |
| Mangrove | 0.914 | 0.819 | 0.878 |  | 0.062 | 0.254 |
| Freshwater swamp | 0.956 | 0.891 | 0.932 | 0.878 |  | 0.093 |
| Coastal forest | 0.908 | 0.704 | 0.940 | 0.658 | 0.756 |  |

### Relationship between insect and plant richness

Compared to mangroves (ca. 250 plant species), rainforest and swamp forest sites have 4.6 or 7.6 times the number of recorded plant species based on checklists for the sites (Table S5). This higher species richness is also confirmed by plot data for the rainforest [69] (839 species in 52 plots of 100m^2^) and swamp forest [70] (671 species in 40 plots of 400m^2^). However, the insect biodiversity of the rainforest and swamp forest sites is only 1.64 – 1.98 times higher than in the mangroves after rarefaction (1.99 – 2.52 times higher in the core dataset).

### Analysis of ecological guilds and correlation between insect and plant diversity

For this analysis, we used the core dataset consisting of Diptera and Hymenoptera which dominate Malaise trap samples (see Brown 2005 [71] & Hebert et al. 2016 [72]). The chosen taxa occupy a broad range of ecological guilds. Also note that this core dataset consists of specimens from traps with at least 6 months continuous sampling from April – September in order to include one dry and wet season. In order to understand how different habitats maintain insect diversity, we assigned insect species with known family/genus identities to ecological guilds (42,092 specimens belonging to 2,230 putative species; no guild information is available for the remaining 19,974 specimens) After stepwise refinement of an analysis of covariance (ANCOVA) model, the final model was defined as: *insectdiv ~ habitat + guild + plantdiv + guild:plantdiv* (*insectdiv*: rarefied insect species richness, *plantdiv*: plant species richness). The type-II sum of squares test reveal that guild and the interaction term between guild and plant diversity are highly significant factors (p < 0.001), while plant diversity (p = 0.063) and habitat (p = 0.468) are not. This suggests guild and plant diversity together play an important role in determining insect diversity but the precise relationship warranted further testing. Single variable linear regressions (*insectdiv ~ plantdiv)* were performed on each guild separately (Fig. 3) and plant diversity was found to only be highly significantly and positively correlated with the alpha-diversity of phytophagous and fungivorous insects (p < 0.001, R^2^ = 0.992 and 0.990, p = 0.886 and 0.943 respectively).

**Figure 3.**
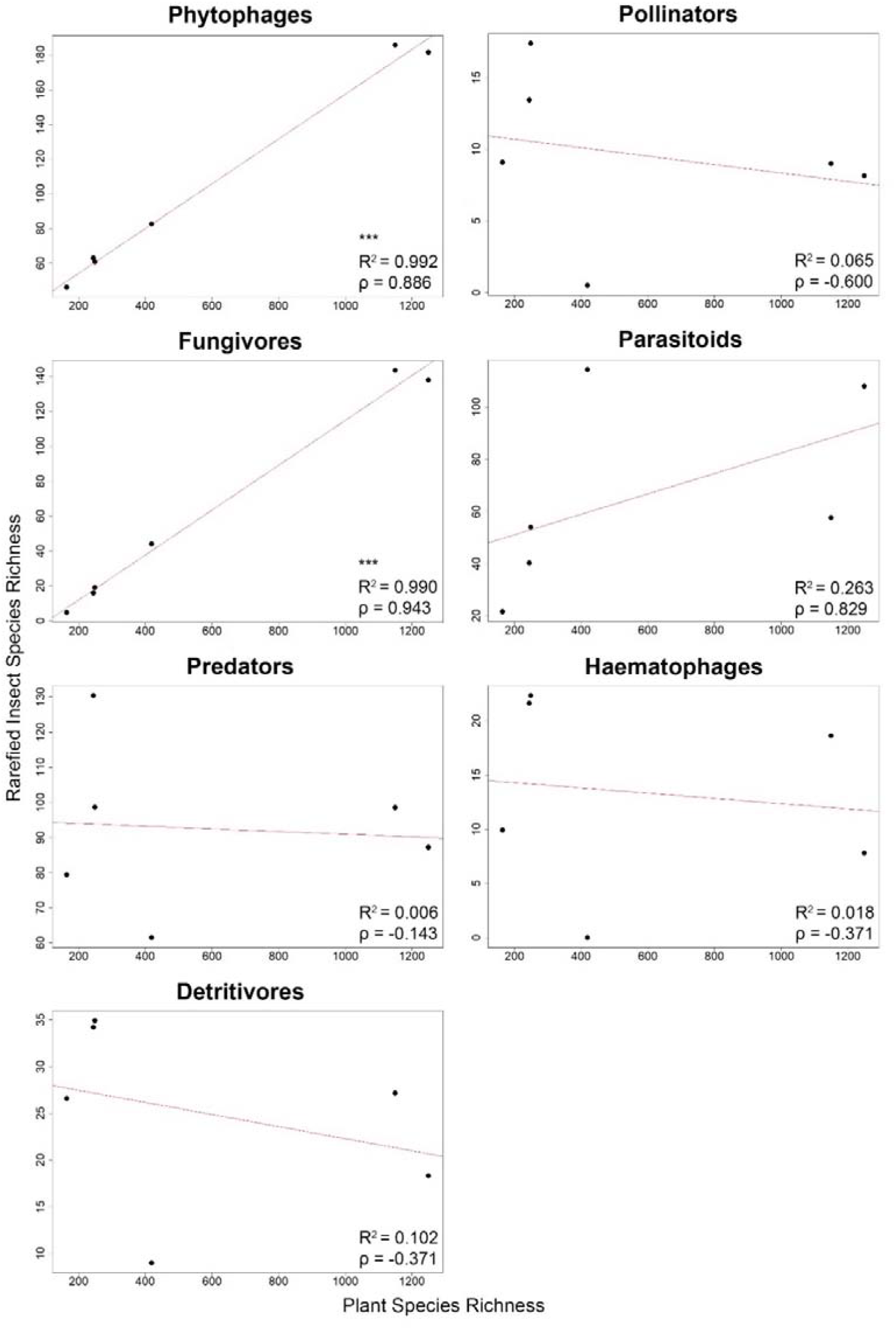
Only the diversity of phytophagous and fungivorous insects is correlated with plant diversity based on a linear regression model using rarefied insect species richness (*: ≤0.05, **: ≤0.01, ***: ≤0.001).

The different habitat types vary in composition (Fig. 4 & Table S11). Rainforest and freshwater swamp forest sites have higher numbers and proportions of phytophagous and fungivorous insect species (see also Figs S7 & S8). The insect communities of mangroves, however, are characterized by an elevated proportion of predatory species. With regard to species turnover, communities are separated by habitat for most guilds and pairwise comparisons (Fig. 5, Tables S12 & S13).

**Figure 4.**
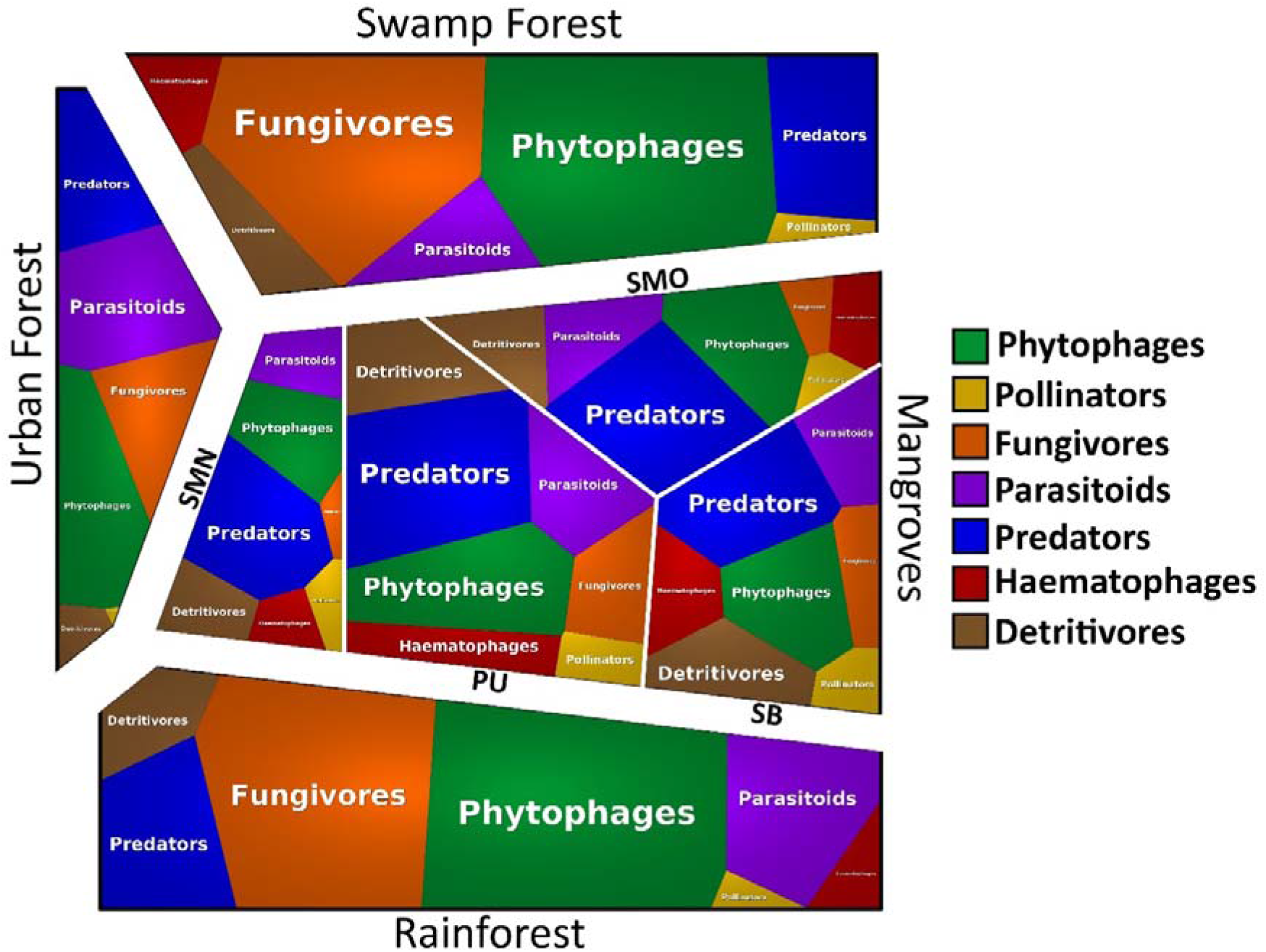
Voronoi treemap of insect guilds across four habitats. Phytophages and fungivores dominate in rain and swamp forests while predators are overrepresented in mangroves. Mangroves are represented by four sites (PU=Pulau Ubin, SB=Sungei Buloh, SMO: Semakau old mangrove, SMN: Semakau restored mangrove).

**Figure 5.**
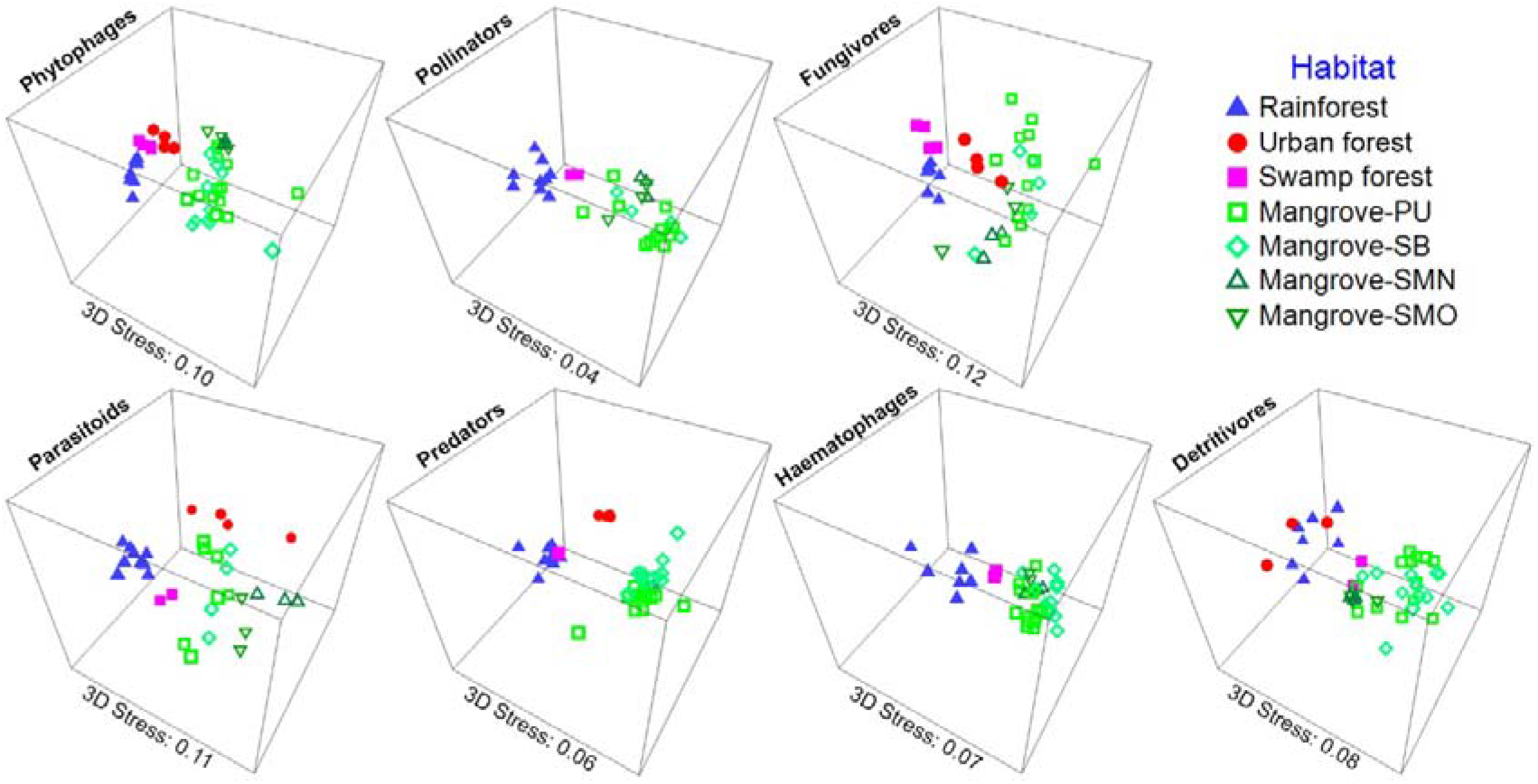
Habitat differentiation by insect guilds (3D NMDS plot of Bray-Curtis distances for habitats with >2 sites).

### Species turnover across Asian mangroves

The specimens from Hong Kong belonged to 109 dolichopodid, 129 phorid, and 25 mycetophilid species. The corresponding number for Brunei were 96 and 76 species for dolichopodids and phorids, with too few mycetophilids being available for evaluation (Table S4). The southern Thai dolichopodids belonged to 74 species. We find high species turnover between Hong Kong, Brunei and Singapore, even after rarefying the specimen sample sizes (Fig. S9). Approximately 90% of all dolichopodid and phorid species are unique to each region and <1% are shared across all regions. Species turnover is even higher for the mycetophilids of Hong Kong and Singapore (>95%). Species turnover for the dolichopodids of Southern Thailand and Singapore is again high with only 11.5% of all species shared between both countries.

## Discussion

### Discovery of a largely overlooked, predator-enriched insect community in mangroves

It is often assumed that the insect diversity in mangroves is low because high salinity and low plant diversity are thought to be unfavourable to insect diversification [23, 73, 74]. However, we here show that mangroves are species-rich for insects despite low plant diversity (<250 species: [75–77]). This result is unambiguously supported by rarefaction, but also supported by species richness estimation with frequency ratio and CNE estimators although the latter are likely to overestimate the species richness because the estimates triple the observed species richness and the variances are high [66]. In addition to being species-rich, the mangrove fauna is also very unique. More than half of its species are not found in other habitats, even though coastal forests are adjacent to mangroves. Indeed, after adjusting for sampling effort, the species diversity in Singapore’s premier rainforest reserve (Bukit Timah Nature Reserve: 1.64 km^2^) and largest swamp forest remnant (Nee Soon: 5 km^2^) is only 50% higher than the diversity of major mangrove sites (PU: 0.904 km^2^, SB: 1.168 km^2^, SM: 0.174 km^2^). The high diversity encountered in the mangrove sites was particularly unexpected because the rainforests of Bukit Timah Nature Reserve have been protected for more than 50 years [78, 79] and have very high plant diversity (e.g., 1,250 species of vascular plants [69] including 341 species of trees [80] in a 2 ha plot of the Centre for Tropical Forest Science). Moreover, we extensively sampled the insect diversity in the reserve by placing multiple Malaise traps in primary, maturing secondary, and old secondary forests. Similarly, we expected the insect diversity of Singapore’s largest swamp forest (Nee Soon) to greatly exceed the number of species found in the mangrove sites because the swamp forest is also known for its high species richness (e.g., 1,150 species of vascular plant species [81]).

A guild-level analysis reveals that rainforests and swamp forests have overall the highest species diversity for most guilds (Fig. S7). Mangroves maintain high species diversity although they are impoverished for phytophagous and fungivorous species. They are, however, home to a disproportionally large number of predatory species (Fig. 4) whose larvae develop in sediments (Empidoidea and Tabanidae). This suggests that the high insect diversity in tropical habitats may be achieved by having large proportions of species developing in the biologically most productive microhabitats – plants and fungi for many forest habitats and the highly productive mud flats for mangroves.

In addition to finding high alpha-diversity in mangroves, we also document that the mangrove insect communities are very distinct. This conclusion is supported by a multitude of analyses (mvabund, ANOSIM, SIMPER, NMDS). It is furthermore insensitive to the removal of rare species (Fig. 2) and driven by high species turnover rather than nestedness (see Table 1C). This stratification by habitat is still evident even when the two dominant insect orders in Malaise trap samples (Diptera and Hymenoptera) are removed (Fig. 2). Comparatively high overlap is only observed between mangroves and coastal forests (860 shared species) which is presumably due to the close proximity of the habitats on Pulau Ubin (Fig. S1) where back mangroves and coastal forests are contiguous. The uniqueness of the mangrove insect community is probably due to the unusual environmental conditions characterized by extreme daily fluctuations in salinity, temperature, and inundation. These extreme conditions are likely to require physiological and behavioural adaptations that encourage the emergence of an evolutionarily distinct fauna. What is surprising, however, is that we find no evidence for an adaptive radiation of particular clades. Instead, a large number of independent colonization events seems more likely given that the mangrove species usually belong to genera that are also known from other habitats (e.g., Dolichopodidae). This challenges the view that high salinity is a potent colonization barrier for invertebrates [73, 74].

Mangrove regeneration is pursued in many countries, with mixed success in restoring the original plant diversity [82, 83], but it remains poorly understood whether regenerated mangroves harbour the original arthropod biodiversity. Our preliminary data based on 311 Malaise trap samples from one regenerated site suggests that this is not the case. The regenerated mangrove (SMN) was replanted with a monoculture of *Rhizophora stylosa* [77] which replaced old-growth mangroves that had been cleared during reclamation work (1994–1999 [51]). The restored site (SMN) has markedly lower insect species richness than all other mangrove sites, including a neighbouring old-growth mangrove (SMO; Fig. 1). This highlights once more that habitat assessments have to be holistic and should not only be based on plant and vertebrate data [84].

Mangrove insect communities are not only rich and distinct in Singapore. Within Asia, we reveal a 92% species turnover between Singapore and Hong Kong (2,500 km north; Fig. S1) for taxa representing different guilds (Dolichopodidae–predators: 483 species, Mycetophilidae–fungivores: 67 species, Phoridae–mostly saprophagous: 591 species). While climatic differences could be advanced as a potential explanation, comparisons with the mangroves in the geographically close and tropical Borneo (Brunei) confirm a high species turnover of 85% (see also Grootaert 2019 [85]). Further evidence for high regional species turnover in mangroves emerges when the dolichopodid fauna of Singapore’s and Brunei’s mangroves are compared with the fauna of Southern Thailand (coasts of South China and Andaman seas). Only 34 and 10 of the 74 known Thai species are shared with Singapore and Brunei respectively; These data suggest that a significant proportion of the global insect diversity may reside in mangroves. Based on the data from Singapore, it appears that much of the diversity may still be intact, given that we find no evidence that the insect diversity in Singapore’s mangrove fragments is depressed relative to what is found in the much more pristine sites in Brunei. This also suggests that the loss of species diversity for small, flying insects in Singapore may not have been as dramatic as what has been documented for Singapore’s vertebrate and large invertebrate species [48, 86, 87].

### Discovering a new insect hotspot with NGS barcoding

Global insect declines have recently received much attention by the scientific community [2] and public [88]. Obtaining relevant data is very difficult since quantifying insect diversity using conventional techniques is slow and expensive. This is because too many specimens have to be sorted into too many species before a holistic habitat assessment can be carried out [89]. In our study, this problem is overcome by specimen sorting using NGS barcodes, which differ from traditional barcodes by costing only a fraction of those obtained with Sanger sequencing. Based on previous tests, we find that species delimited with 313-bp NGS barcodes have 90% congruence with species-level units delimited with morphological data [56, 57, 90, 91]. This suggests that large-scale species discovery with NGS barcodes can yield sufficiently accurate information on species abundance and distribution for habitat assessments [55, 56]. This also means that NGS barcodes could be used for quickly revealing hidden hotspots of insect diversity in countries with high diversity and limited science funding. We estimate that the ~140,000 specimens in our study could today be sequenced for <USD25,000 using 350 manpower days whereas a similar study based on morphology would require >150 manpower years [92]; i.e. some of the traditional obstacles to understanding arthropod biodiversity caused by the taxonomic impediment may be finally disappearing. However, it is important to remember that barcode-derived units are only proxies for formal species and should only be used for broad analyses of diversity patterns. Additional taxonomic work may uncover, for example, rapid radiations in particular habitats or guilds that would be overlooked by studies that rely on clustering barcodes based on similarity [93]. This is why the rich dolichopodid fauna is currently being revised using the “reverse workflow” approach where morphological validation follows presorting with barcodes [56, 85, 94].

### Concluding remarks

We here document that the insect fauna inhabiting mangroves is not only rich, but also distinct when compared to many other tropical forest habitats. The discovery of such an unexpectedly rich and distinct insect community highlights how little we know about arthropod diversity. We predict that advances in sequencing technology will facilitate the discovery of numerous additional insect diversity hotspots in tropical and temperate habitats. Mangroves will likely be only one of many future additions to the growing list of habitats that have only recently been recognized as biodiversity hotspots (e.g., dry forests [95, 96], forest savannahs [97, 98]). Our study highlights that accelerating species discovery is a pressing task given that many of these habitats are disappearing at a much faster rate than tropical rainforests and coral reefs.

## Methods

### Sampling site, sample collection, and processing

Singapore has a large number of tropical habitat types that are all within 40 km of each other without being separated by major physical barriers. This allowed us to sample rainforests (from early secondary to mature secondary forest), urban-edge forests, mangroves, swamp forests, freshwater swamps and dry coastal forests. The freshwater swamp habitat differs from swamp forests by largely lacking tree-cover, while the dry coastal forests are distinct from the mangroves by lacking typical mangrove tree species. Note that the habitats had experienced similar levels of habitat degradation or loss due to urbanization (>95% loss of original vegetation cover [48]; *ca*. 90% loss for rainforests [49]; *ca*. 93% loss of swamp forest [50]; 91% loss for mangroves [51]). We sampled these habitat types using 107 trapping sites (Fig. S1). The mangrove sites were located primarily along the North-western and Southern coasts of the mainland, as well as on offshore islands in the south and northeast. The major mangrove sites were on Pulau Ubin (PU), Sungei Buloh (SB) and Pulau Semakau (SM), the last of which is represented by an old-growth (SMO) and a newly regenerated mangrove fragment (SMN).The swamp forest site (Nee Soon) was Singapore’s largest remaining freshwater swamp remnant which is known for a rich insect fauna [99], overall high species richness, and level of endemism [100, 101]. Bukit Timah Nature Reserve was selected as the tropical rainforest site given its high species diversity and protected status [78]. This reserve consists of forests in various stages of succession and hence we sampled different forest types with three sites each being in primary forest, old secondary forest, and maturing secondary forest. The “urban secondary forest” sites were located along a disturbance gradient ranging from the campus of the National University of Singapore (NUS) through several urban parks and forest edges in Central and South Singapore. The freshwater swamp site is located primarily in Kranji, a freshwater marsh at the flooded edge of a reservoir. The “coastal forest” sites were dry secondary forests adjacent to the coast at Labrador Park and Pulau Ubin, which are also close to urban settlements.

All specimens were collected between 2012–2019 (Table S1) using Malaise traps. These traps are widely used for insect surveys because they are effective sampling tools for flying insects and allow for standardized, long-term sampling. Note that the use of Malaise traps in our study was appropriate because the canopy height was comparable for most habitats given that we compared mature mangroves (PU, SB and SMO) with a wet swamp forest site, and different kinds of secondary forests (pers. obs.). Only the canopy height of some sites in Bukit Timah Nature Reserve (BTNR) was higher, but for BTNR we also included secondary forests and several traps were placed on steep slopes that would be able to sample canopy-active fauna from a lower elevation. With regard to the habitat patches, the fragments were larger for the rainforest and swamp forest than for any of the mangrove sites (tropical rainforest: 1.64 km^2^; swamp forest: 5 km^2^, mangrove forest fragments: 0.904 km^2^ [PU], 1.168 km^2^ [SB], 0.174 km^2^ [SM] [51]). Malaise traps in the mangroves were set up in the intertidal zone. Each Malaise trap sample consisted of one-week’s worth of insects preserved in molecular grade ethanol.

After an ethanol change, the specimens were sorted by para-taxonomists (Table S2) into mostly orders or families, but in some cases, we used broader categories that could be communicated to the parataxonomists but did not represent formal taxa: Dermaptera, Lepidoptera, Mantodea, Neuroptera, Orthoptera, Psocodea and Trichoptera, Arachnida (subsampled into Araneae and mites), Blattodea (subsampled into cockroaches, termites), Hemiptera (subsampled into Reduviidae, Dipsocoromorpha, other Heteroptera, Auchenorrhyncha, Sternorrhyncha, other Homoptera), Hymenoptera (subsampled into: Apoidea, Formicidae), Coleoptera (subsampled into Cicindelinae, Mordellidae, Staphylinidae, Elateridae, Cantharidae, Buprestidae, Curculionoidea, other Coleoptera), and Diptera (subsampled into Tipuloidea, Mycetophilidae, Keroplatidae, Culicidae, Chironomidae, Sciaridae, other Nematocera, Dolichopodidae, other Empidoidea, Stratiomyidae, Syrphidae, Asilidae, Tabanidae, Phoridae, other Acalyptrates, Calyptrates), All subsequent barcoding used these subsamples as sampling units. Overall, the samples were typical for Malaise traps in that they were dominated by Diptera and Hymenoptera (>75% of specimens: Table S2, see Fig. S2 for species composition).

We therefore first developed a core dataset for these orders given that the specimen numbers were sufficiently high (62,066 specimens, 4,002 species at 3% p-distance). This core dataset consisted of 12 Diptera families, “other Acalyptrates”, ants (Formicidae) and Apoidea (see Table 2). It also only included specimens from sites that were sampled from 2012 – 2015 and overlapped with regard to the April – September period which covers one dry and one wet season. We subsequently added the available data for other sites and orders. For most of these, we barcoded all available specimens (Araneae, Blattodea, Dermaptera, Mantodea, Neuroptera, Orthoptera, Trichoptera). Three insect orders had intermediate abundances (Coleoptera, Lepidoptera, Hemiptera, Psocodea). For Coleoptera, we processed easily recognized families like the low-abundance insect orders; i.e., all specimens collected before 2019 were sequenced (Staphylinidae, Cleridae, Cerambycidae, Scirtidae, Carabidae, and Elateridae). For Lepidoptera, we processed all large specimens and only sequenced micromoths. For Hemiptera and Psocodea, we barcoded all Heteroptera, but only subsampled the psocodean and homopteran planthoppers because the abundances were too high (planthoppers: 6987 specimens across 6 habitats; Psocodea: 539 specimens across 4 habitats). Here, the subsampling was similar to what had been done for the core data set in that we barcoded all specimens for the samples of particular time periods.

After obtaining barcodes, the taxa in the core dataset were re-identified to genus or family based on morphological re-examination or DNA barcodes that were submitted to the Global Biodiversity Information Facility (GBIF: www.gbif.org) or the Barcode of Life Data (BOLD: www.boldsystems.org) databases. Only matches above 95% and 97% similarity were considered sufficiently precise for family- and genus-level matches respectively. In some cases, these identifications revealed sorting errors by the parataxonomists. The barcodes were nevertheless kept for the diversity analyses because such sorting errors are random across samples.

The mangrove specimens from Hong Kong were collected by 24 Malaise traps installed between October 2017 to October 2018, while those from Brunei were collected by six Malaise traps from July to November 2014. Dolichopodidae, Phoridae, and Mycetophilidae were pre-sorted and send for barcoding to Singapore. Note that the mangrove forests in Brunei are less affected by urbanization than those in Singapore. The dolichopodid specimens from Thailand were obtained by different techniques including sweep-netting from 42 mangrove sites over a period of 15 months from Mar 2014 – Dec 2015.

### Putative species sorting with NGS barcoding

NGS barcoding combines the advantages of cost-effective sequencing with Illumina with the approximate species-level resolution provided by DNA barcodes. The molecular procedures can be learned in hours and several hundred specimens can be processed per person and day. The overall barcode costs are now <10 cents per specimen if Illumina Novaseq is used for sequencing (2 cents/barcode based on USD 6,900 per 250-bp PE flow cell yielding 800 million reads: https://research.ncsu.edu/gsl/pricing). We used NGS barcoding to amplify and sequence a 313-bp fragment of the cytochrome oxidase I gene (*cox1*) using a protocol described in Meier et al. [55]. Direct-PCR [102] was conducted for specimens collected early in the study; during this phase, we used 1-2 legs of the specimen as template for obtaining the amplicon with the primer pair mlCO1intF: 5’-GGWACWGGWTGAACWGTWTAYCCYCC-3’ [103] and jgHCO2198: 5’-TANACYTCNGGRTGNCCRAARAAYCA-3’ [104]. For samples processed later, the whole specimen was immersed in Lucigen QuickExtract solution or HotSHOT buffer [105] and gDNA extraction was conducted non-destructively. The gDNA extract was then used as a PCR template with the afore-mentioned reagents and protocol. The primers used were labelled with 9-bp long barcodes that differed by at least three base pairs. Every specimen in each sequencing library was assigned a unique combination of labelled forward and reverse primers, which allowed the Illumina reads to be binned according to specimen. A negative control was prepared and sequenced for each 96-well PCR plate. Amplification success rates for each plate were assessed via gel electrophoresis for eight random wells per plate.

The amplicons were pooled at equal volumes within each plate and later pooled across plates. Equimolarity was estimated by the presence and intensity of bands on gels. The pooled samples were cleaned with Bioline SureClean Plus and/or via gel cuts before outsourcing library preparation to AITbiotech using TruSeq Nano DNA Library Preparation Kits (Illumina) or the Genome Institute of Singapore (GIS) using NEBNext DNA Library Preparation Kits (NEB). Paired-end sequencing was performed on Illumina Miseq (2×300-bp or 2×250-bp) or Hiseq 2500 platforms (2×250-bp) over multiple runs, thereby allowing troubleshooting and re-sequencing for specimens which initially failed to yield a sufficiently large number of reads. Some of the specimens were also sequenced on the MinION (Oxford Nanopore) platform using primers with a slightly longer tags (13-bp) and following the protocol described in Srivathsan et al. [106, 57]. Raw Illumina reads were processed with the bioinformatics pipeline and quality-control filters described in Meier et al. [55]. A BLAST search to GenBank’s nucleotide (nt) database was also conducted to identify and discard contaminants by parsing the BLAST output through *readsidentifier* [107] and removing barcodes with incorrect matches at >97% identity.

To obtain putative species units, the *cox1* barcodes were clustered over a range of uncorrected p-distance thresholds (2–4%) typically used for species delimitation in the literature [108]. The clustering was performed with a python script that implements the objective clustering algorithm of Meier et al. 2006 [59] and allows for large scale processing. USEARCH [60] (cluster_fast) was used to confirm the results by setting -id at 0.96, 0.97 and 0.98. To gauge how many of our species/specimens matched barcodes in public databases, we used the “Sequence ID” search of the Global Biodiversity Information Facility (GBIF). We then determined the number of matches with identity scores <97. We then counted the number of matches to barcodes with species-level identifications.

### Diversity analyses

For analysis of species richness and turnover, we initially focused on a core dataset of Diptera and Hymenoptera (62,066 specimens, 4,002 species at 3% p-distance) that consisted of several taxa representing a range of ecological guilds (see Table 2). This dataset included the sites that were sampled from 2012 – 2015 and overlap in the April – September period, in order to control for seasonal and sampling effects. We subsequently analysed the full dataset consisting of all barcodes obtained in the project, which included more taxa and trapping sites. Results for the full dataset are described here because they are broadly congruent with those for the core dataset. The latter are in the supplementary. To assess the species richness of the six major habitat types, samples were rarefied with the *iNEXT* [109] R package (R Development Core Team) using 1,000 bootstrap replicates in order to account for unequal sampling completeness. The rarefaction was performed by coverage [61] in the main analysis (Fig. 1) and by specimen count in the supplementary (Fig. S3). Site comparisons were carried out by comparing species diversity post-rarefaction to the lowest coverage/smallest number of specimens. The habitat type “mangrove” was treated both as a single habitat as well as separate sites (PU, SB, SMN, SMO, others) in separate analyses. We also obtained species richness extrapolations for each habitat using a frequency ratio estimator [62] and a combined non-parametric estimator (CNE) [63] based on combined Chao1 and Chao2 extrapolations. Both were performed in R, with the former using the *breakaway* package and the latter using R code published with the original manuscript.

In order to study species turnover, we first excluded 11 trapping sites that had <100 specimens in total so as to prevent poor sampling from inflating site distinctness. We then determined the distinctness of the communities across habitats using non-metric multidimensional scaling (NMDS) plots that were prepared with PRIMER v7 [110] using Bray-Curtis dissimilarity. Plots were generated for each habitat type and the different mangrove sites; Bray-Curtis was chosen because it is a preferred choice for datasets that include abundance information. The dataset was split into three groups: the dominant orders (Diptera and Hymenoptera) and all others combined, in order to test if the results were driven by the dominant orders. Community structuring by habitat was verified by multivariate analysis of variance (MANOVA) tests performed in R with the *manova* function, with habitat as the explanatory variable and the response variable being combined NMDS1 and NMDS2 coordinates, as well as both coordinates separately. The NMDS1 and NMDS2 coordinates were derived with the *metaMDS* function from *vegan* in R [111]. We also employ a model-based framework for testing if habitat influenced abundance distribution by coding the former as the explanatory variable and the latter as the response variable in *mvabund* [67] in R. The *manyglm* function was used to construct the model, with negative binomial error distribution. Lastly, analysis of similarities (ANOSIM) and similarity percentages (SIMPER) were performed in PRIMER under default parameters in order to obtain ANOSIM p-values and R-statistics for both the entire dataset and the pairwise comparisons between habitat types. The SIMPER values were calculated for within and between-habitat types. The ANOSIM p-values can be used to assess significant differences while the R-statistic allows for determining the degree of similarity, with values closer to 1 indicating greater distinctness. We also used the *betapart* [112] R package to examine if the observed dissimilarity (Bray-Curtis) was due to species turnover or nestedness. The *beta*.*multi*.*abund* and *beta*.*pair*.*abund* functions were used to split the global and pairwise dissimilarity scores into turnover and nestedness components. Lastly, the robustness of the results was tested by removing singleton, doubleton and rare species (<5 and <10 individuals) from the datasets. The pruned datasets were subjected to the same analyses as the full dataset. For the guild-specific datasets, traps with fewer than three species were excluded in the species turnover analyses because large distances driven by undersampling can obscure signal.

To examine species turnover across larger geographic scales, dolichopodid, phorid, and mycetophilid specimens from Singapore were compared with those from Hong Kong (Dolichopodidae: 2,601; Phoridae: 562, Mycetophilidae: 186), and Brunei (Dolichopodidae: 2,800; Phoridae: 272), and data for the dolichopodids of Southern Thai mangroves (942 specimens). Since Singapore was more extensively sampled, the Singaporean dataset was randomly subsampled (10 iterations in Microsoft Excel with the RAND() function) to the number of specimens available for the other two countries (Table S4). The species diversity after rarefaction was then compared (with 95% confidence intervals for the rarefied data).

### Ecological guild and plant diversity analyses

For the guild-level analysis, we focused on the core dataset consisting of the species belonging to the two dominant orders Diptera and Hymenoptera. This dataset consists of 62,066 specimens from 9 rainforest, 4 swamp forest, 4 urban forest, and 32 mangrove sites (Fig. S1). In order to test for an overall correlation between plant and insect diversity, we obtained data for the plant diversity in the respective sites from checklists and survey plots (Table S5). In order to further examine the correlation between plant and insect diversity across multiple ecological guilds, we assigned the identified Diptera and Hymenoptera families and genera non-exclusively to ecological guilds (phytophages, pollinators, fungivores, parasitoids, predators, haematophages and detritivores) based on known adult and larval natural history traits for the group (Table S3). Taxa with different adult and larval natural histories are placed in both guilds. Taxa lacking sufficient information or with highly variable life-history strategies were assigned to the “Others/Unknown” category and excluded from analysis.

Barcodes from each guild were separately aligned and clustered at 3% p-distance. For taxa that have adults and immatures with different natural histories (i.e., belong to two distinct ecological guilds), the species counts were halved and placed into both guilds when calculating rarefied species abundance and richness. Guild distribution for each habitat was visualized with Voronoi treemaps via Proteomaps [113]. Species turnover for the guild-specific subsets were analysed with PRIMER to generate NMDS plots, as well as ANOSIM and SIMPER values. The rarefied species richness values were also used for a multivariate model analysis. An ANCOVA model was constructed in R [114] with the *lm* function: *insectdiv ~ site * habitat * guild * plantdiv*, with insectdiv representing rarefied insect alpha-diversity and plantdiv representing plant species counts. The “site” factor was excluded due to collinearity and the model was refined via stepwise removal of factors starting with the most complex (interaction terms) and least significant ones. At each stage, the *anova* function was used to assess loss of informational content and the final model was derived when the reported p-value was significant (p < 0.05). The model’s residuals were examined to ensure the data were normal. Subsequently, the *Anova* function from the *car* package [115] was used to obtain type-II test statistics. Finally, single-variable linear regression was performed in R with the *lm* function: *insectdiv ~ plantdiv* for each guild separately to obtain significance, multiple R-squared and Spearman’s rho values.

## Acknowledgements

All work described here was carried out as part of a comprehensive insect survey of Singapore which was carried out in collaboration and with support from the National Parks Board of Singapore (NParks). Special thanks go to the team from the National Biodiversity Centre of NParks for their assistance in fieldwork (Permits: NP/RP12-022-4, NP/RP12-022-5, NP/RP12-022-6) We would also like to thank the research staff, lab technicians, undergraduate students and interns of the Evolutionary Biology Laboratory for their help and assistance. This project would have been impossible without their hard work. Special thanks go to Lee Wan Ting, Yuen Huei Khee and Arina Adom. Financial support was provided by a Ministry of Education grant on biodiversity discovery (R-154-000-A22-112). The Hong Kong Mangroves project is supported by the Environment and Conservation Fund (ECF Project 69/2016) and we thank Dr. Christopher Taylor, Mr. Roy Shun-Chi Leung, and Ms. Ukyoung Chang, for their help in taking and sorting the samples and Dr Stefano Cannicci for his lead in the mangrove project. Mangrove insects in Brunei were sampled with permission from Brunei Forestry Department and Ministry of Primary Resources and Tourism during an UBD postdoctoral fellowship awarded to Claas Damken (Research and collecting permit file numbers: UBD/CAN–387(b)(SAA); UBD/ADM/R3(z)Pt.; UBD/PNC2/2/RG/1(293)). We thank Roman Carrasco for commenting on the manuscript.

## Availability of Data and Materials

The 313bp cox1 sequence data analysed in this paper will be made public on acceptance.

Reviewers may access these data via this private link: https://figshare.com/s/1381706292a55cfba13b

## Author information

*National University of Singapore, Department of Biological Sciences*

Darren Yeo, Amrita Srivathsan, Jayanthi Puniamoorthy & Rudolf Meier

*Lee Kong Chian Natural History Museum, Singapore*

Foo Maosheng & Ang Yuchen

*Royal Belgian Institute of Natural Sciences, Brussels, Belgium and National Biodiversity Centre of National Parks Board*

Patrick Grootaert

*The University of Hong Kong, School of Biological Sciences*

Benoit Guénard

*Universiti Brunei Darussalam, Institute for Biodiversity and Environmental Research*

Claas Damken & Rodzay A. Wahab

*National Parks Board, Singapore, International Biodiversity Conservation Division*

Lena Chan

## Author contributions

Darren Yeo: design of the work, acquisition, analysis, and interpretation of data, drafting and revising manuscript

Amrita Srivathsan: creation of new software used in the work, analysis and interpretation of data, revision of manuscript

Jayanthi Puniamoorthy: design of the work, acquisition of data, revision of manuscript

Foo Maosheng: design of the work, acquisition of data, revision of manuscript

Patrick Grootaert: conception and design of the work, acquisition of data, revision of manuscript

Lena Chan: conception and design of the work, revision of manuscript Benoit Guénard: conception (Hong Kong), revision of manuscript

Claas Damken: conception (Brunei), acquisition of data, revision of manuscript Rodzay A. Wahab: conception (Brunei), reading of manuscript

Ang Yuchen: identification of Diptera taxa, revision of manuscript

Rudolf Meier: conception and design of the work, data analysis and interpretation, drafted and revised manuscript

## Competing interests

None

## Notes

### Competing Interest Statement

The authors have declared no competing interest.

### Summary of Updates

text and figure changes

## References

1. Dirzo R, Young HS, Galetti M, Ceballos G, Isaac NJB, Collen B. Defaunation in the Anthropocene. Science. 2014;345:401–6.

2. Hallmann CA, Sorg M, Jongejans E, Siepel H, Hofland N, Schwan H, et al. More than 75 percent decline over 27 years in total flying insect biomass in protected areas. PLoS ONE. 2017;12:e0185809.

3. Lister BC, Garcia A. Climate-driven declines in arthropod abundance restructure a rainforest food web. Proc Natl Acad Sci USA. 2018;115:E10397–406.

4. Díaz S, Settele J, Brondízio E, Ngo HT, Guèze M, Agard J, et al. Summary for policymakers of the global assessment report on biodiversity and ecosystem services of the Intergovernmental Science-Policy Platform on Biodiversity and Ecosystem Services. IPBES. 2019;:45.

5. Basset Y, Lamarre GPA. Toward a world that values insects. Science. 2019;364:1230–1.

6. Cardoso P, Branco VV, Chichorro F, Fukushima CS, Macías-Hernández N. Can we really predict a catastrophic worldwide decline of entomofauna and its drivers? Global Ecology and Conservation. 2019;20:e00621.

7. Komonen A, Halme P, Kotiaho JS. Alarmist by bad design: Strongly popularized unsubstantiated claims undermine credibility of conservation science. ReEco. 2019;4:17–9.

8. Sánchez-Bayo F, Wyckhuys KAG. Worldwide decline of the entomofauna: A review of its drivers. Biological Conservation. 2019;232:8–27.

9. Simmons BI, Balmford A, Bladon AJ, Christie AP, De Palma A, Dicks LV, et al. Worldwide insect declines: An important message, but interpret with caution. Ecol Evol. 2019;9:3678–80.

10. Thomas CD, Jones TH, Hartley SE. “Insectageddon”: A call for more robust data and rigorous analyses. Glob Change Biol. 2019;25:1891–2.

11. Coddington JA, Agnarsson I, Miller JA, Kuntner M, Hormiga G. Undersampling bias: the null hypothesis for singleton species in tropical arthropod surveys. Journal of Animal Ecology. 2009;78:573–84.

12. Stork NE. How many species of insects and other terrestrial arthropods are there on Earth? Annu Rev Entomol. 2018;63:31–45.

13. Basset Y, Cizek L, Cuénoud P, Didham RK, Novotny V, Ødegaard F, et al. Arthropod distribution in a tropical rainforest: Tackling a four dimensional puzzle. PLoS ONE. 2015;10:e0144110.

14. Kishimoto-Yamada K, Itioka T. How much have we learned about seasonality in tropical insect abundance since Wolda (1988)? Entomological Science. 2015;18:407–19.

15. van Klink R, Bowler DE, Gongalsky KB, Swengel AB, Gentile A, Chase JM. Meta-analysis reveals declines in terrestrial but increases in freshwater insect abundances. Science. 2020;368:417–20.

16. Seastedt TR, Crossley DA. The Influence of Arthropods on Ecosystems. BioScience. 1984;34:157– 61.

17. Perfecto I, Vandermeer J, Hanson P, Cartiân V. Arthropod biodiversity loss and the transformation of a tropical agro-ecosystem. Biodiversity and Conservation. 1997;6:935–45.

18. Whiles MR, Charlton RE. The ecological significance of tallgrass prairie arthropods. Annu Rev Entomol. 2006;51:387–412.

19. Ewers RM, Boyle MJW, Gleave RA, Plowman NS, Benedick S, Bernard H, et al. Logging cuts the functional importance of invertebrates in tropical rainforest. Nat Commun. 2015;6:6836.

20. Bar-On YM, Phillips R, Milo R. The biomass distribution on Earth. Proc Natl Acad Sci USA. 2018;115:6506–11.

21. Barlow J, França F, Gardner TA, Hicks CC, Lennox GD, Berenguer E, et al. The future of hyperdiverse tropical ecosystems. Nature. 2018;559:517–26.

22. Novotny V, Basset Y. Rare species in communities of tropical insect herbivores: pondering the mystery of singletons. Oikos. 2000;89:564–72.

23. Novotny V, Drozd P, Miller SE, Kulfan M, Janda M, Basset Y, et al. Why are there so many species of herbivorous insects in tropical rainforests? Science. 2006;313:1115–8.

24. Basset Y, Cizek L, Cuenoud P, Didham RK, Guilhaumon F, Missa O, et al. Arthropod Diversity in a Tropical Forest. Science. 2012;338:1481–4.

25. Duke NC, Meynecke J-O, Dittmann S, Ellison AM, Anger K, Berger U, et al. A world without mangroves? Science. 2007;317:41b–2b.

26. Valiela I, Bowen JL, York JK. Mangrove forests: One of the world’s threatened major tropical environments. BioScience. 2001;51:807.

27. Gilman EL, Ellison J, Duke NC, Field C. Threats to mangroves from climate change and adaptation options: A review. Aquatic Botany. 2008;89:237–50.

28. Spencer T, Schuerch M, Nicholls RJ, Hinkel J, Lincke D, Vafeidis AT, et al. Global coastal wetland change under sea-level rise and related stresses: The DIVA Wetland Change Model. Global and Planetary Change. 2016;139:15–30.

29. Friess DA, Richards DR, Phang VXH. Mangrove forests store high densities of carbon across the tropical urban landscape of Singapore. Urban Ecosyst. 2016;19:795–810.

30. Alongi DM. Mangrove forests: Resilience, protection from tsunamis, and responses to global climate change. Estuarine, Coastal and Shelf Science. 2008;76:1–13.

31. Spalding MD, Ruffo S, Lacambra C, Meliane I, Hale LZ, Shepard CC, et al. The role of ecosystems in coastal protection: Adapting to climate change and coastal hazards. Ocean & Coastal Management. 2014;90:50–7.

32. Zavalloni M, Groeneveld RA, van Zwieten PAM. The role of spatial information in the preservation of the shrimp nursery function of mangroves: A spatially explicit bio-economic model for the assessment of land use trade-offs. Journal of Environmental Management. 2014;143:17–25.

33. Nagelkerken I, Blaber SJM, Bouillon S, Green P, Haywood M, Kirton LG, et al. The habitat function of mangroves for terrestrial and marine fauna: A review. Aquatic Botany. 2008;89:155–85.

34. Yates KK, Rogers CS, Herlan JJ, Brooks GR, Smiley NA, Larson RA. Diverse coral communities in mangrove habitats suggest a novel refuge from climate change. Biogeosciences. 2014;11:4321–37.

35. Zhang K, Lin S, Ji Y, Yang C, Wang X, Yang C, et al. Plant diversity accurately predicts insect diversity in two tropical landscapes. Mol Ecol. 2016;25:4407–19.

36. Bagchi R, Gallery RE, Gripenberg S, Gurr SJ, Narayan L, Addis CE, et al. Pathogens and insect herbivores drive rainforest plant diversity and composition. Nature. 2014;506:85–8.

37. Becerra JX. On the factors that promote the diversity of herbivorous insects and plants in tropical forests. Proc Natl Acad Sci USA. 2015;112:6098–103.

38. Moreira X, Abdala-Roberts L, Rasmann S, Castagneyrol B, Mooney KA. Plant diversity effects on insect herbivores and their natural enemies: current thinking, recent findings, and future directions. Current Opinion in Insect Science. 2016;14:1–7.

39. Lewinsohn TM, Novotny V, Basset Y. Insects on Plants: Diversity of Herbivore Assemblages Revisited. Annu Rev Ecol Evol Syst. 2005;36:597–620.

40. Batista-da-Silva JA. Effect of lunar phases, tides, and wind speed on the abundance of Diptera Calliphoridae in a mangrove swamp. Neotrop Entomol. 2014;43:48–52.

41. Chowdhury S. Butterflies of Sundarban Biosphere Reserve, West Bengal, eastern India: a preliminary survey of their taxonomic diversity, ecology and their conservation. J Threat Taxa. 2014;6:6082–92.

42. Rohde C, Silva DMIO, Oliveira GF, Monteiro LS, Montes MA, Garcia ACL. Richness and abundance of the cardini group of Drosophila (Diptera, Drosophilidae) in the Caatinga and Atlantic Forest biomes in northeastern Brazil. An Acad Bras Ciênc. 2014;86:1711–8.

43. Hazra AK, Dey MK, Mandal GP. Diversity and distribution of arthropod fauna in relation to mangrove vegetation on a newly emerged island on the river Hooghly, West Bengal. Rec Zool Surv India. 2005;104:99–102.

44. Adeduntan SA, Olusola JA. Diversity and abundance of arthropods and tree species as influenced by different forest vegetation types in Ondo State, Nigeria. International Journal of Ecosystem. 2013;3:19–23.

45. García-Gómez A, Castaño-Meneses G, Vázquez-González MM, Palacios-Vargas JG. Mesofaunal arthropod diversity in shrub mangrove litter of Cozumel Island, Quintana Roo, México. Applied Soil Ecology. 2014;83:44–50.

46. D’Cunha P, Nair VM. Diversity and distribution of ant fauna in Hejamadi Kodi sandspit, Udupi district, Karnataka, India. Halteres. 2013;4:33–47.

47. Balakrishnan S, Srinivasan M, Mohanraj J. Diversity of some insect fauna in different coastal habitats of Tamil Nadu, southeast coast of India. Journal of Asia-Pacific Biodiversity. 2014;7:408–14.

48. Brook BW, Sodhi NS, Ng PKL. Catastrophic extinctions follow deforestation in Singapore. Nature. 2003;424:420–3.

49. Corlett RT. Plant succession on degraded land in Singapore. Journal of Tropical Forest Science. 1991;4:151–61.

50. Davison GWH, Cai Y, Li TJ, Lim WH. Integrated research, conservation and management of Nee Soon freshwater swamp forest, Singapore: hydrology and biodiversity. GBS 70(Suppl1) Nee Soon. 2018;70:1–7.

51. Yee ATK, Ang WF, Teo S, Liew SC, Tan HTW. The present extent of mangrove forests in Singapore. Nature in Singapore. 2010;3:139–45.

52. Colwell RK, Coddington J. Estimating terrestrial biodiversity through extrapolation. Phil Trans R Soc Lond B. 1994;345:101–18.

53. Longino JT, Coddington J, Colwell RK. The ant fauna of a tropical rain forest: estimating species richness three different ways. Ecology. 2002;83:689–702.

54. Scharff N, Coddington JA, Griswold CE, Hormiga G, Bjørn P de P. When to quit? estimating spider species richness in a northern european deciduous forest. Journal of Arachnology. 2003;31:246–73.

55. Meier R, Wong W, Srivathsan A, Foo M. $1 DNA barcodes for reconstructing complex phenomes and finding rare species in specimen-rich samples. Cladistics. 2016;32:100–10.

56. Wang WY, Srivathsan A, Foo M, Yamane SK, Meier R. Sorting specimen-rich invertebrate samples with cost-effective NGS barcodes: Validating a reverse workflow for specimen processing. Mol Ecol Resour. 2018;18:490–501.

57. Srivathsan A, Hartop E, Puniamoorthy J, Lee WT, Kutty SN, Kurina O, et al. Rapid, large-scale species discovery in hyperdiverse taxa using 1D MinION sequencing. preprint. Evolutionary Biology; 2019. doi:10.1101/622365.

58. Yeo D, Srivathsan A, Meier R. Longer is not always better: Optimizing barcode length for large-scale species discovery and identification. preprint. Evolutionary Biology; 2019. doi:10.1101/594952.

59. Meier R, Shiyang K, Vaidya G, Ng PKL. DNA barcoding and taxonomy in Diptera: A tale of high intraspecific variability and low identification success. Systematic Biology. 2006;55:715–28.

60. Edgar RC. Search and clustering orders of magnitude faster than BLAST. Bioinformatics. 2010;26:2460–1.

61. Chao A, Jost L. Coverage-based rarefaction and extrapolation: standardizing samples by completeness rather than size. Ecology. 2012;93:2533–47.

62. Willis A, Bunge J. Estimating diversity via frequency ratios: Estimating Diversity via Ratios. Biom. 2015;71:1042–9.

63. Ronquist F, Forshage M, Häggqvist S, Karlsson D, Hovmöller R, Bergsten J, et al. Completing Linnaeus’s inventory of the Swedish insect fauna: Only 5,000 species left? PLoS ONE. 2020;15:e0228561.

64. Chiu C-H, Chao A. Estimating and comparing microbial diversity in the presence of sequencing errors. PeerJ. 2016;4:e1634.

65. Mahé F, de Vargas C, Bass D, Czech L, Stamatakis A, Lara E, et al. Parasites dominate hyperdiverse soil protist communities in Neotropical rainforests. Nat Ecol Evol. 2017;1:0091.

66. Gotelli NJ, Colwell RK. Estimating species richness. Biological diversity: frontiers in measurement and assessment. 2011;12:39–54.

67. Wang Y, Naumann U, Wright ST, Warton DI. mvabund - an R package for model-based analysis of multivariate abundance data: The mvabund R package. Methods in Ecology and Evolution. 2012;3:471–4.

68. Benjamin DJ, Berger JO, Johannesson M, Nosek BA, Wagenmakers E-J, Berk R, et al. Redefine statistical significance. Nat Hum Behav. 2018;2:6–10.

69. Ho BC, Lua HK, Ibrahim B, Yeo RSW, Athen P, Leong PKF, et al. The plant diversity in Bukit Timah Nature Reserve, Singapore. GBS. 2019;71 suppl.1:41–144.

70. Chong KY, Lim RCJ, Loh JW, Neo L, Seah WW, Tan SY, et al. Rediscoveries, new records, and the floristic value of the Nee Soon freshwater swamp forest, Singapore. GBS 70(Suppl1) Nee Soon. 2018;70:49–69.

71. Brown BV. Malaise Trap Catches and the Crisis in Neotropical Dipterology. American Entomologist. 2005;51:180–3.

72. Hebert PDN, Ratnasingham S, Zakharov EV, Telfer AC, Levesque-Beaudin V, Milton MA, et al. Counting animal species with DNA barcodes: Canadian insects. Phil Trans R Soc B. 2016;371:20150333.

73. Berezina NA. Tolerance of Freshwater Invertebrates to Changes in Water Salinity. Russian Journal of Ecology. 2003;34:261–6.

74. Silberbush A, Blaustein L, Margalith Y. Influence of Salinity Concentration on Aquatic Insect Community Structure: A Mesocosm Experiment in the Dead Sea Basin Region. Hydrobiologia. 2005;548:1–10.

75. Lee S, Leong’ SAP, Ibrahim’ A, Gwee’ A-T. A Botanical Survey of Chek Jawa, Pulau Ubin, Singapore. Gardens’ Bulletin Singapore. 2003;55:271–307.

76. Tan HTW, Choong MF, Chua KS, Loo AHB, bin Haji Ahmad HS, Seah EEL, et al. A botanical survey of Sungei Buloh Nature Park, Singapore. The Gardens’ Bulletin. 1997;49:15–35.

77. Teo S, Yeo RKH, Chong KY, Chung YF, Neo L, Tan HTW. The flora of Pulau Semakau: A Project Semakau checklist. Nature in Singapore. 2011;4:263–72.

78. Corlett RT. Bukit Timah: the History and Significance of a Small Rain-forest Reserve. Envir Conserv. 1988;15:37–44.

79. Chan L, Davison GWH. Synthesis of results from the Comprehensive Biodiversity Survey of Bukit Timah Nature Reserve, Singapore, with recommendations for management. GBS. 2019;71 suppl.1:583–610.

80. LaKrankie J, Davies SJ, Wang LK, Lee SK, Lum SKY. Forest Trees of Bukit Timah. Population ecology in a tropical forest fragment. Singapore: Simply Green; 2005.

81. Wong HF, Tan SY, Koh CY, Siow HJM, Li T, Heyzer A, et al. Checklist of the plant species of Nee Soon swamp forest, Singapore: Bryophytes to Angiosperms. National Parks Board and Raffles Museum of Biodiversity Research, National University of Singapore, Singapore. 2013.

82. Lee SY, Hamilton S, Barbier EB, Primavera J, Lewis RR. Better restoration policies are needed to conserve mangrove ecosystems. Nat Ecol Evol. 2019;3:870–2.

83. Kodikara KAS, Mukherjee N, Jayatissa LP, Dahdouh-Guebas F, Koedam N. Have mangrove restoration projects worked? An in-depth study in Sri Lanka: Evaluation of mangrove restoration in Sri Lanka. Restor Ecol. 2017;25:705–16.

84. Chua MAH. The herpetofauna and mammals of Semakau landfill: A Project Semakau checklist. Nature in Singapore. 2011;4:277–87.

85. Grootaert P. Species turnover between the northern and southern part of the South China Sea in the Elaphropeza Macquart mangrove fly communities of Hong Kong and Singapore (Insecta: Diptera: Hybotidae). EJT. 2019. doi:10.5852/ejt.2019.554.

86. Lane DJW, Kingston T, Lee BPY-H. Dramatic decline in bat species richness in Singapore, with implications for Southeast Asia. Biological Conservation. 2006;131:584–93.

87. Sodhi NS, Wilcove DS, Lee TM, Sekercioglu CH, Subaraj R, Bernard H, et al. Deforestation and Avian Extinction on Tropical Landbridge Islands: Bird Extinction on Tropical Islands. Conservation Biology. 2010;24:1290–8.

88. Mikanowski J. ‘A different dimension of loss’: inside the great insect die-off. The Guardian. 2017. https://www.theguardian.com/environment/2017/dec/14/a-different-dimension-of-loss-great-insect-die-off-sixth-extinction. Accessed 20 Sep 2019.

89. Novotny V, Miller SE. Mapping and understanding the diversity of insects in the tropics: past achievements and future directions: Mapping tropical diversity of insects. Austral Entomology. 2014;53:259–67.

90. Grootaert P. Revision of the genus Thinophilus Wahlberg (Diptera: Dolichopodidae) from Singapore and adjacent regions: A long term study with a prudent reconciliation of a genetic to a classic morphological approach. Raffles Bulletin of Zoology. 2018;66:413–73.

91. Yeo D, Puniamoorthy J, Ngiam RWJ, Meier R. Towards holomorphology in entomology: rapid and cost-effective adult-larva matching using NGS barcodes: Life-history stage matching with NGS barcodes. Syst Entomol. 2018;43:678–91.

92. Basset Y, Cizek L, Cuénoud P, Didham RK, Guilhaumon F, Missa O, et al. How many arthropod species live in a tropical forest. Science. 2012;338:1481–4.

93. Meier R, Blaimer B, Buenaventura E, Hartop E, von Rintelen T, Srivathsan A, et al. A re-analysis of the data in Sharkey et al.’s (2021) minimalist revision reveals that BINs do not deserve names, but BOLD Systems needs a stronger commitment to open science. preprint. Evolutionary Biology; 2021. doi:10.1101/2021.04.28.441626.

94. Hartop E, Srivathsan A, Ronquist F, Meier R. Large-scale Integrative Taxonomy (LIT): resolving the data conundrum for dark taxa. preprint. Evolutionary Biology; 2021. doi:10.1101/2021.04.13.439467.

95. Janzen DH. Insect diversity of a Costa Rican dry forest: why keep it, and how? Biological Journal of the Linnean Society. 1987;30:343–56.

96. Banda-R K, Delgado-Salinas A, Dexter KG, Linares-Palomino R, Oliveira-Filho A, Prado D, et al. Plant diversity patterns in neotropical dry forests and their conservation implications. Science. 2016;353:1383.

97. Beirão MV, Neves FS, Penz CM, DeVries PJ, Fernandes GW. High butterfly beta diversity between Brazilian cerrado and cerrado–caatinga transition zones. J Insect Conserv. 2017;21:849–60.

98. Pennington RT, Lehmann CER, Rowland LM. Tropical savannas and dry forests. Current Biology. 2018;28:R541–5.

99. Baloğlu B, Clews E, Meier R. NGS barcoding reveals high resistance of a hyperdiverse chironomid (Diptera) swamp fauna against invasion from adjacent freshwater reservoirs. Front Zool. 2018;15:31.

100. Ng PKL, Lim KKP. The conservation status of the Nee Soon freshwater swamp forest of Singapore. Aquatic Conserv: Mar Freshw Ecosyst. 1992;2:255–66.

101. Turner IM, Min BC, Kwan WY, Ting CP. Freshwater swamp forest in Singapore, with particular reference to that found around the Nee Soon firing ranges. The Gardens’ Bulletin, Singapore. 1996;48:129–57.

102. Wong WH, Tay YC, Puniamoorthy J, Balke M, Cranston PS, Meier R. ‘Direct PCR’ optimization yields a rapid, cost-effective, nondestructive and efficient method for obtaining DNA barcodes without DNA extraction. Mol Ecol Resour. 2014;14:1271–80.

103. Leray M, Yang JY, Meyer CP, Mills SC, Agudelo N, Ranwez V, et al. A new versatile primer set targeting a short fragment of the mitochondrial COI region for metabarcoding metazoan diversity: application for characterizing coral reef fish gut contents. Front Zool. 2013;10:34.

104. Geller J, Meyer C, Parker M, Hawk H. Redesign of PCR primers for mitochondrial cytochrome c oxidase subunit I for marine invertebrates and application in all-taxa biotic surveys. Mol Ecol Resour. 2013;13:851–61.

105. Montero-Pau J, Gómez A, Muñoz J. Application of an inexpensive and high-throughput genomic DNA extraction method for the molecular ecology of zooplanktonic diapausing eggs: Rapid DNA extraction for diapausing eggs. Limnol Oceanogr Methods. 2008;6:218–22.

106. Srivathsan A, Baloğlu B, Wang W, Tan WX, Bertrand D, Ng AHQ, et al. A MinION™-based pipeline for fast and cost-effective DNA barcoding. Mol Ecol Resour. 2018;18:1035–49.

107. Srivathsan A, Sha JCM, Vogler AP, Meier R. Comparing the effectiveness of metagenomics and metabarcoding for diet analysis of a leaf-feeding monkey (Pygathrix nemaeus). Mol Ecol Resour. 2015;15:250–61.

108. Ratnasingham S, Hebert PDN. A DNA-based registry for all animal species: The Barcode Index Number (BIN) system. PLoS ONE. 2013;8:e66213.

109. Chao A, Gotelli NJ, Hsieh TC, Sander EL, Ma KH, Colwell RK, et al. Rarefaction and extrapolation with Hill numbers: a framework for sampling and estimation in species diversity studies. Ecological Monographs. 2014;84:45–67.

110. Clarke KR, Gorley RN. Primer. PRIMER-e, Plymouth. 2006.

111. Oksanen J, Blanchet FG, Friendly M, Kindt R, Legendre P, McGlinn D, et al. vegan: Community Ecology Package. R package version 25–7. 2020.

112. Baselga A, Orme CDL. betapart⍰: an R package for the study of beta diversity: Betapart package. Methods in Ecology and Evolution. 2012;3:808–12.

113. Liebermeister W, Noor E, Flamholz A, Davidi D, Bernhardt J, Milo R. Visual account of protein investment in cellular functions. Proceedings of the National Academy of Sciences. 2014;111:8488– 93.

114. R Development Core Team. R: a language and environment for statistical computing. R Foundation for Statistical Computing, Vienna, Austria, www.R-project.org. 2005.

115. Fox J, Weisberg S. An R companion to applied regression. Sage publications; 2018.

116. Tan HTW, Lim KKP, Tan MK, Yee ATK. The Biology of Kent Ridge: Vegetation, Plants and Animals. In: Kent Ridge: An Untold Story. Singapore: NUS Press; 2019. p. 49–172.

