## Supplementary material for "Mangroves are an overlooked hotspot of insect diversity despite low plant diversity": Supplmentary Materials

**Table S1.** Collection periods and trap localities; M = mangroves, SF = swamp forest, UF = urban forest, TF = tropical rainforest, CF = coastal forest, FS = freshwater swamp. Numbers in brackets indicate traps with <100 specimens in total that were excluded from the diversity analyses.

| Sampling period | Location | Habitat type | GPS coordinates | No. of traps | Total no. of weekly samples | Used for guild-level analyses |
| --- | --- | --- | --- | --- | --- | --- |
| Singapore | | | | | | |
| Apr 2012 – Mar 2014 | Pulau Ubin | M | 1°24'36.3"N 103°59'25.5"E | 3 | 72 | Y |
|  | Pulau Semakau original | M | 1°12'17.6"N 103°45'37.7"E | 3 | 72 | Y |
|  | Pulau Semakau replanted | M | 1°12'03.1"N 103°45'45.4"E | 3 | 72 | Y |
|  | Sungei Buloh Wetland Reserve | M | 1°26'46.3"N 103°43'49.9"E | 2 | 48 | Y |
|  | Nee Soon freshwater swamp | SF | 1°23'00.3"N 103°48'46.5"E | 2 | 48 | Y |
| May 2014 – Jun 2014 | Mandai Nature Park | M | 1°26'18.3"N 103°45'49.7"E | 3 (1) | 6 | N |
|  | Pulau Tekong | M | 1°25'47.3"N 104°03'46.3"E | 3 (2) | 6 | N |
|  | Sarimbun | M | 1°25'59.1"N 103°41'21.8"E | 3 (1) | 6 | N |
| Nov 2014 – May 2015 | Nee Soon freshwater swamp | SF | 1°23'00.3"N 103°48'46.5"E | 2 | 14 | Y |
| Apr 2015 – Sep 2015 | NUS | UF | 1°17'49.6"N 103°46'35.7"E | 4 | 24 | Y |
| Mar 2016 – Aug 2016 | Pulau Ubin | M | 1°25'11.64"N 103°56'6.25"E | 10 | 60 | Y |
|  | Sungei Buloh Wetland Reserve | M | 1°26'43.20"N 103°43'5.10"E | 10 | 60 | Y |
|  | Labrador Park | M | 1°16'13.3"N 103°48'10.1"E | 3 | 18 | N |
|  | Labrador Park | CF | 1°16'05.4"N 103°48'16.2"E | 2 | 18 | N |
| Aug 2016 – Oct 2017 | Bukit Timah Nature Reserve primary forest | TF | 1°21'13.90"N 103°46'47.57"E | 3 | 45 | Y |
|  | Bukit Timah Nature Reserve old secondary forest | TF | 1°21'17.96"N 103°46'54.01"E | 3 | 45 | Y |
|  | Bukit Timah Nature Reserve maturing secondary forest | TF | 1°21'4.57"N 103°46'53.80"E | 3 | 45 | Y |
| Apr 2017 – 20 Sep 2017 | NUS | UF | 1°17'49.6"N 103°46'35.7"E | 4 | 18 | N |
| Sep 2017 – Dec 2017 | NUS | UF | 1°17'45.3"N 103°46'13.8"E | 3 (1) | 16 | N |
| Mar 2018 – Jun 2018 | Bishan-Ang Moh Kio Park | UF | 1°21'35.7"N 103°50'49.9"E | 2 | 26 | N |
|  | Enabling Village | UF | 1°17'13.6"N 103°48'53.3"E | 1 | 13 | N |
|  | Esplanade Theatre | UF | 1°17'26.4"N 103°51'17.9"E | 1 | 13 | N |
|  | Sungei Buloh Wetland Reserve | M | 1°26'52.45"N 103°43'24.16"E | 4 (1) | 16 | N |
|  | Kranji Marshes | FS | 1°25'0.56"N 103°43'43.50"E | 3 | 12 | N |
|  | Lim Chu Kang | M | 1°26'48.80"N 103°42'35.71"E | 2 | 8 | N |
|  | Mandai Nature Park | M | 1°26'37.96"N 103°45'59.70"E | 4 | 16 | N |
|  | Pulau Ubin | CF | 1°24'26.3"N 103°57'16.3"E | 3 | 12 | N |
|  | Pulau Ubin | M | 1°24'32.2"N 103°57'12.1"E | 8 | 32 | N |
|  | Labrador Park | M | 1°16'13.3"N 103°48'10.1"E | 5 | 20 | N |
|  | Labrador Park | CF | 1°16'05.4"N 103°48'16.2"E | 4 (2) | 20 | N |
| Mar 2019 – Jun 2019 | Coney Island | M | 1°24'37.3"N 103°55'23.1"E | 5 (2) | 15 | N |
|  | Kranji Marshes | FS | 1°25'11.0"N 103°43'54.3"E | 4 | 15 | N |
|  | Pulau Ubin | CF | 1°25'34.7"N 103°56'29.2"E | 1 | 15 | N |
|  | Pulau Ubin | M | 1°25'05.3"N 103°56'06.5"E | 7 (6) | 15 | N |
| Hong Kong | | | | | | |
| Nov 2017 – Dec 2017,  May 2018 – Jul 2018 | Ha Pak Nai | M | 22°25'31.48"N 113°56'20.11"E | 6 | 30 | Y |
|  | Hang Mei | M | 22°15'9.83"N 113°52'5.84"E | 5 | 25 | Y |
|  | Ho Chung | M | 22°21'13.18"N 114°15'7.45"E | 6 | 30 | Y |
|  | Lai Chi Wo | M | 22°31'37.63"N 114°15'43.63"E | 5 | 25 | Y |
|  | Nam Chung | M | 22°31'31.62"N 114°12'28.94"E | 5 | 25 | Y |
|  | Sai Keng | M | 22°25'13.48"N 114°16'4.66"E | 5 | 25 | Y |
|  | Sam A Chung | M | 22°30'29.84"N 114°16'20.93"E | 5 | 25 | Y |
|  | Sam A Tsuen | M | 22°30'55.22"N 114°16'16.36"E | 5 | 25 | Y |
|  | Sha Tau Kok | M | 22°32'4.34"N 114°12'39.78"E | 10 | 50 | Y |
|  | Sheung Pak Nai | M | 22°27'7.09"N 113°57'45.11"E | 5 | 25 | Y |
|  | Shui Hau | M | 22°13'9.70"N 113°55'8.33"E | 5 | 25 | Y |
|  | So Lo Pun | M | 22°32'17.20"N 114°15'21.49"E | 5 | 25 | Y |
|  | Tai O | M | 22°15'28.44"N 113°51'48.96"E | 6 | 30 | Y |
|  | Tai Tam | M | 22°14'46.10"N 114°13'24.02"E | 3 | 15 | Y |
|  | Tai Tan | M | 22°26'18.85"N 114°19'59.77"E | 1 | 5 | Y |
|  | To Kwa Peng | M | 22°25'43.07"N 114°19'59.30"E | 5 | 25 | Y |
|  | Tsim Bei Tsui | M | 22°29'20.47"N 113°59'53.95"E | 5 | 25 | Y |
|  | Tung Chung | M | 22°16'52.50"N 113°55'44.04"E | 6 | 30 | Y |
|  | Wong Chuk Wan | M | 22°23'44.27"N 114°17'10.21"E | 5 | 25 | Y |
|  | Yim Tin Tsai | M | 22°22'32.74"N 114°18'5.76"E | 5 | 25 | Y |
| Brunei | | | | | | |
| Jul 2014 – Nov 2014 | Pulau Berambang | M | 4°54'7.44"N 115°1'17.94"E | 2 | 10 | Y |
|  | Labu Forest Reserve | M | 4°51'41.75"N 115°6'59.69"E | 2 | 10 | Y |
|  | Tutong Forest | M | 4°46'9.54"N 114°36'20.64"E | 2 | 10 | Y |

**Table S2.** Specimen counts from the full dataset (core dataset shaded in green) with taxa split by habitat type and site (for mangroves only).

| Order | Family | Coastal Forest | Freshwater Swamp | Rainforest | Swamp Forest | Urban Forest | Mangrove PU | Mangrove SB | Mangrove SMN | Mangrove SMO | Mangrove Others |
| --- | --- | --- | --- | --- | --- | --- | --- | --- | --- | --- | --- |
| Araneae |  | 45 | 2 | 409 | 188 | 981 | 1080 | 782 | 320 | 544 | 172 |
| Blattodea | **Termitoidae** | 38 | 0 | 977 | 80 | 83 | 649 | 74 | 0 | 12 | 4 |
|  | **Other Blattodea** | 10 | 0 | 183 | 152 | 71 | 88 | 57 | 14 | 54 | 7 |
| Coleoptera |  | 54 | 26 | 915 | 574 | 0 | 92 | 78 | 57 | 265 | 2 |
| Dermaptera |  | 0 | 0 | 1 | 0 | 0 | 0 | 0 | 0 | 0 | 0 |
| Hemiptera |  | 76 | 10 | 2617 | 365 | 816 | 979 | 501 | 606 | 768 | 246 |
| Lepidoptera |  | 513 | 53 | 619 | 0 | 1 | 159 | 13 | 16 | 103 | 17 |
| Mantodea |  | 27 | 0 | 99 | 2 | 35 | 13 | 10 | 2 | 7 | 0 |
| Neuroptera | **Coniopterygidae** | 0 | 0 | 23 | 0 | 0 | 2 | 0 | 0 | 0 | 0 |
|  | **Other Neuroptera** | 0 | 0 | 1 | 0 | 0 | 0 | 0 | 0 | 0 | 0 |
| Orthoptera |  | 9 | 3 | 56 | 27 | 33 | 180 | 214 | 109 | 208 | 0 |
| Psocoptera |  | 141 | 15 | 283 | 0 | 0 | 97 | 2 | 0 | 0 | 1 |
| Trichoptera |  | 0 | 0 | 1 | 0 | 0 | 0 | 0 | 0 | 0 | 0 |
| Hymenoptera | **Formicidae** | 258 | 372 | 189 | 1055 | 3329 | 1406 | 292 | 789 | 730 | 85 |
|  | **Apoidea** | 302 | 289 | 2130 | 58 | 353 | 151 | 416 | 146 | 211 | 56 |
| Diptera | **Asilidae** | 14 | 2 | 29 | 12 | 9 | 105 | 33 | 45 | 73 | 41 |
|  | **Culicidae** | 226 | 3913 | 89 | 338 | 2 | 738 | 613 | 336 | 597 | 95 |
|  | **Dolichopodidae** | 1078 | 898 | 743 | 262 | 4143 | 7966 | 1945 | 7841 | 4070 | 1210 |
|  | **Empididae** | 77 | 7 | 41 | 106 | 69 | 472 | 18 | 1 | 0 | 3 |
|  | **Hybotidae** | 1 | 7 | 0 | 6 | 0 | 10 | 3 | 6 | 1 | 1 |
|  | **Keroplatidae** | 56 | 4 | 5 | 137 | 48 | 18 | 10 | 12 | 90 | 0 |
|  | **Mycetophilidae** | 202 | 272 | 874 | 1176 | 650 | 82 | 121 | 10 | 52 | 23 |
|  | **Phoridae** | 2406 | 2178 | 2122 | 345 | 6955 | 2242 | 808 | 293 | 302 | 308 |
|  | **Stratiomyidae** | 53 | 29 | 66 | 80 | 212 | 333 | 241 | 148 | 101 | 80 |
|  | **Syrphidae** | 34 | 28 | 19 | 10 | 6 | 150 | 141 | 14 | 36 | 43 |
|  | **Tabanidae** | 51 | 164 | 27 | 43 | 0 | 467 | 113 | 59 | 66 | 66 |
|  | **Tephritidae** | 4 | 0 | 42 | 21 | 8 | 397 | 70 | 652 | 607 | 13 |
|  | **Other Brachycera** | 1249 | 5840 | 1192 | 3574 | 2353 | 5444 | 1812 | 3148 | 3359 | 3485 |
|  | **Other Nematocera** | 2179 | 7894 | 1917 | 865 | 145 | 2154 | 46 | 34 | 65 | 403 |

**Table S3.** Diptera and Hymenoptera species used in the guild-level analyses are identified to higher taxonomic levels where possible and assigned to ecological guild based on known natural history traits (grey = guild assignment; some taxa are listed for several guilds because immatures and adults have different known natural histories).

| **Taxon** | | **Ecological Guild** | | | | | | | | | | | |
| --- | --- | --- | --- | --- | --- | --- | --- | --- | --- | --- | --- | --- | --- |
| **Family** | **Genus** | **Phytophages** | **Pollinators** | **Fungivores** | | **Parasitoids** | | **Predators** | **Haematophages** | | **Detritivores** | **Others/Unknown** | |
| **Diptera** | | | | | | | | | | | | | |
| **Agromyzidae** |  | ⌧ |  | |  | |  |  |  |  | | |  |
| **Anthomyiidae** |  |  |  | |  | |  |  |  |  | | | ⌧ |
| **Asilidae** |  |  |  | |  | |  | ⌧ |  |  | | |  |
| **Asteiidae** |  |  |  | | ⌧ | |  |  |  |  | | | ⌧ |
| **Athericidae** |  |  |  | |  | |  | ⌧ |  |  | | | ⌧ |
| **Bombyliidae** |  |  | ⌧ | |  | |  | ⌧ |  |  | | |  |
| **Calliphoridae** |  |  |  | |  | |  |  |  |  | | | ⌧ |
| **Canacidae** |  | ⌧ |  | |  | |  |  |  |  | | |  |
| **Chloropidae** | *Anacamptoneurum* | ⌧ |  | |  | |  |  |  |  | | |  |
| **Chloropidae** | *Cadrema* |  |  | |  | |  |  |  | ⌧ | | |  |
| **Chloropidae** | *Chlorops* | ⌧ |  | |  | |  |  |  |  | | |  |
| **Chloropidae** | *Chloropsina* | ⌧ |  | |  | |  |  |  |  | | |  |
| **Chloropidae** | *Conioscinella* | ⌧ |  | |  | |  |  |  |  | | |  |
| **Chloropidae** | *Dasyopa* |  |  | |  | |  |  |  |  | | | ⌧ |
| **Chloropidae** | *Gampsocera* |  |  | |  | |  |  |  |  | | | ⌧ |
| **Chloropidae** | *Gaurax* |  |  | |  | |  |  |  |  | | | ⌧ |
| **Chloropidae** | *Lasiambia* |  |  | |  | |  | ⌧ |  |  | | |  |
| **Chloropidae** | *Liohippelates* | ⌧ |  | |  | |  |  |  |  | | |  |
| **Chloropidae** | *Malloewia* |  |  | |  | |  |  |  |  | | | ⌧ |
| **Chloropidae** | *Olcella* | ⌧ |  | |  | |  |  |  |  | | |  |
| **Chloropidae** | *Oscinella* | ⌧ |  | |  | |  |  |  |  | | |  |
| **Chloropidae** | *Polyodaspis* |  |  | |  | |  | ⌧ |  |  | | |  |
| **Chloropidae** | *Pseudogaurax* |  |  | |  | |  | ⌧ |  |  | | |  |
| **Chloropidae** | *Pseudopachychaeta* | ⌧ |  | |  | |  |  |  |  | | |  |
| **Chloropidae** | *Rhodesiella* |  |  | |  | |  |  |  | ⌧ | | |  |
| **Chloropidae** | *Thaumatomyia* |  |  | |  | |  | ⌧ |  |  | | |  |
| **Chloropidae** | *Thyridula* |  |  | |  | |  |  |  |  | | | ⌧ |
| **Chloropidae** | *Tricimba* |  |  | |  | |  |  |  | ⌧ | | |  |
| **Clusiidae** |  | ⌧ |  | |  | |  |  |  |  | | | ⌧ |
| **Coelopidae** |  | ⌧ |  | |  | |  |  |  |  | | |  |
| **Cryptochetidae** |  |  |  | |  | | ⌧ |  |  |  | | |  |
| **Culicidae** |  |  |  | |  | |  |  | ⌧ | ⌧ | | |  |
| **Diastatidae** |  |  |  | |  | |  |  |  | ⌧ | | |  |
| **Diopsidae** |  | ⌧ |  | |  | |  |  |  | ⌧ | | |  |
| **Dolichopodidae** |  |  |  | |  | |  | ⌧ |  |  | | |  |
| **Drosophilidae** | *Apenthecia* | ⌧ |  | |  | |  |  |  |  | | |  |
| **Drosophilidae** | *Chymomyza* | ⌧ |  | |  | |  |  |  |  | | |  |
| **Drosophilidae** | *Colocasiomyia* | ⌧ |  | |  | |  |  |  |  | | |  |
| **Drosophilidae** | *Dichaetophora* |  |  | | ⌧ | |  |  |  |  | | |  |
| **Drosophilidae** | *Drosophila* |  |  | |  | |  |  |  |  | | | ⌧ |
| **Drosophilidae** | *Gitona* | ⌧ |  | |  | |  |  |  |  | | |  |
| **Drosophilidae** | *Hirtodrosophila* |  |  | | ⌧ | |  |  |  |  | | |  |
| **Drosophilidae** | *Hypselothyrea* | ⌧ |  | |  | |  |  |  |  | | |  |
| **Drosophilidae** | *Leucophenga* | ⌧ |  | |  | |  |  |  |  | | |  |
| **Drosophilidae** | *Liodrosophila* | ⌧ |  | |  | |  |  |  |  | | |  |
| **Drosophilidae** | *Luzonimyia* |  |  | |  | |  |  |  |  | | | ⌧ |
| **Drosophilidae** | *Microdrosophila* | ⌧ |  | |  | |  |  |  |  | | |  |
| **Drosophilidae** | *Mycodrosophila* |  |  | | ⌧ | |  |  |  |  | | |  |
| **Drosophilidae** | *Paramycodrosophila* |  |  | | ⌧ | |  |  |  |  | | |  |
| **Drosophilidae** | *Scaptodrosophila* | ⌧ |  | |  | |  |  |  |  | | |  |
| **Drosophilidae** | *Scaptomyza* | ⌧ |  | |  | |  |  |  |  | | |  |
| **Drosophilidae** | *Stegana* | ⌧ |  | |  | |  |  |  |  | | |  |
| **Drosophilidae** | *Zaprionus* | ⌧ |  | |  | |  |  |  |  | | |  |
| **Empididae** |  |  |  | |  | |  | ⌧ |  |  | | |  |
| **Ephydridae** | *Allotrichoma* |  |  | |  | |  |  |  | ⌧ | | |  |
| **Ephydridae** | *Atissa* |  |  | |  | |  |  |  | ⌧ | | |  |
| **Ephydridae** | *Brachydeutera* | ⌧ |  | |  | |  |  |  |  | | |  |
| **Ephydridae** | *Cerobothrium* |  |  | |  | |  |  |  |  | | | ⌧ |
| **Ephydridae** | *Ceropsilopa* | ⌧ |  | |  | |  |  |  |  | | |  |
| **Ephydridae** | *Discocerina* | ⌧ |  | |  | |  |  |  |  | | |  |
| **Ephydridae** | *Donaceus* |  |  | |  | |  |  |  |  | | | ⌧ |
| **Ephydridae** | *Glenanthe* |  |  | |  | |  |  |  |  | | | ⌧ |
| **Ephydridae** | *Hecamedoides* |  |  | |  | |  |  |  |  | | | ⌧ |
| **Ephydridae** | *Hydrellia* | ⌧ |  | |  | |  |  |  |  | | |  |
| **Ephydridae** | *Limnellia* |  |  | |  | |  |  |  |  | | | ⌧ |
| **Ephydridae** | *Nostima* | ⌧ |  | |  | |  |  |  |  | | |  |
| **Ephydridae** | *Notiphila* |  |  | |  | |  |  |  | ⌧ | | |  |
| **Ephydridae** | *Ochthera* |  |  | |  | |  | ⌧ |  |  | | |  |
| **Ephydridae** | *Orasiopa* |  |  | |  | |  |  |  |  | | | ⌧ |
| **Ephydridae** | *Paralimna* | ⌧ |  | |  | |  |  |  |  | | |  |
| **Ephydridae** | *Placopsidella* |  |  | |  | |  | ⌧ |  |  | | |  |
| **Ephydridae** | *Polytrichophora* |  |  | |  | |  |  |  |  | | | ⌧ |
| **Ephydridae** | *Ptilomyia* |  |  | |  | |  |  |  |  | | | ⌧ |
| **Ephydridae** | *Rhynchopsilopa* |  |  | |  | |  |  |  |  | | | ⌧ |
| **Ephydridae** | *Trimerogastra* |  |  | |  | |  |  |  |  | | | ⌧ |
| **Ephydridae** | *Trypetomima* |  |  | |  | |  |  |  |  | | | ⌧ |
| **Ephydridae** | *Zeros* |  |  | |  | |  |  |  |  | | | ⌧ |
| **Hybotidae** |  |  |  | |  | |  | ⌧ |  |  | | |  |
| **Keroplatidae** |  | ⌧ |  | | ⌧ | |  |  |  |  | | |  |
| **Lauxaniidae** |  | ⌧ |  | |  | |  |  |  |  | | |  |
| **Lonchaeidae** |  | ⌧ |  | |  | |  |  |  |  | | |  |
| **Lygistorrhinidae** |  | ⌧ |  | | ⌧ | |  |  |  |  | | |  |
| **Megamerinidae** |  |  |  | |  | |  | ⌧ |  |  | | |  |
| **Micropezidae** |  |  |  | |  | |  |  |  |  | | | ⌧ |
| **Milichiidae** | *Aldrichiomyza* |  |  | |  | |  |  |  |  | | | ⌧ |
| **Milichiidae** | *Leptometopa* |  |  | |  | |  |  |  | ⌧ | | |  |
| **Milichiidae** | *Milichia* |  |  | |  | |  |  |  |  | | | ⌧ |
| **Milichiidae** | *Milichiella* |  |  | |  | |  |  |  | ⌧ | | |  |
| **Milichiidae** | *Neophyllomyza* | ⌧ |  | |  | |  |  |  |  | | |  |
| **Milichiidae** | *Paramyia* |  |  | |  | |  |  |  |  | | | ⌧ |
| **Milichiidae** | *Phyllomyza* |  |  | |  | |  |  |  | ⌧ | | |  |
| **Muscidae** |  |  |  | |  | |  |  |  |  | | | ⌧ |
| **Mycetophilidae** |  | ⌧ |  | | ⌧ | |  |  |  |  | | |  |
| **Neriidae** |  | ⌧ |  | |  | |  |  |  |  | | | ⌧ |
| **Odiniidae** |  |  |  | |  | |  |  |  |  | | | ⌧ |
| **Periscelididae** |  | ⌧ |  | |  | |  |  |  |  | | |  |
| **Phoridae** |  |  |  | |  | |  |  |  |  | | | ⌧ |
| **Pipunculidae** |  |  |  | |  | | ⌧ |  |  |  | | |  |
| **Platypezidae** |  |  |  | | ⌧ | |  |  |  | ⌧ | | |  |
| **Platystomatidae** |  |  |  | |  | |  |  |  |  | | | ⌧ |
| **Psilidae** |  | ⌧ |  | |  | |  |  |  |  | | |  |
| **Pyrgotidae** |  |  |  | |  | | ⌧ |  |  |  | | |  |
| **Rhagionidae** |  |  |  | |  | |  | ⌧ | ⌧ |  | | |  |
| **Rhiniidae** |  |  |  | |  | |  | ⌧ |  |  | | |  |
| **Sarcophagidae** |  |  |  | |  | |  |  |  |  | | | ⌧ |
| **Sciaridae** |  | ⌧ |  | | ⌧ | |  |  |  |  | | |  |
| **Sphaeroceridae** |  |  |  | |  | |  |  |  |  | | | ⌧ |
| **Stratiomyiidae** |  | ⌧ |  | |  | |  |  |  | ⌧ | | |  |
| **Syrphidae** | *Allobaccha* |  | ⌧ | |  | |  | ⌧ |  |  | | |  |
| **Syrphidae** | *Allograpta* |  | ⌧ | |  | |  | ⌧ |  |  | | |  |
| **Syrphidae** | *Asarkina* |  | ⌧ | |  | |  | ⌧ |  |  | | |  |
| **Syrphidae** | *Ceriana* |  | ⌧ | |  | |  |  |  | ⌧ | | |  |
| **Syrphidae** | *Eosmallota* |  | ⌧ | |  | |  |  |  | ⌧ | | |  |
| **Syrphidae** | *Eristalinus* |  | ⌧ | |  | |  |  |  | ⌧ | | |  |
| **Syrphidae** | *Eristalis* |  | ⌧ | |  | |  |  |  | ⌧ | | |  |
| **Syrphidae** | *Eumerus* |  | ⌧ | |  | |  |  |  | ⌧ | | |  |
| **Syrphidae** | *Graptomyza* |  | ⌧ | |  | |  |  |  | ⌧ | | |  |
| **Syrphidae** | *Ischiodon* |  | ⌧ | |  | |  | ⌧ |  |  | | |  |
| **Syrphidae** | *Microdon* |  | ⌧ | |  | |  | ⌧ |  |  | | |  |
| **Syrphidae** | *Paragus* |  | ⌧ | |  | |  | ⌧ |  |  | | |  |
| **Syrphidae** | *Psilota* |  | ⌧ | |  | |  |  |  | ⌧ | | |  |
| **Syrphidae** | *Spheginobaccha* |  | ⌧ | |  | |  |  |  |  | | | ⌧ |
| **Syrphidae** | *Syritta* |  | ⌧ | |  | |  |  |  | ⌧ | | |  |
| **Syrphidae** | *Volucella* |  | ⌧ | |  | |  |  |  | ⌧ | | |  |
| **Tabanidae** |  |  |  | |  | |  | ⌧ | ⌧ |  | | |  |
| **Tachinidae** |  |  |  | |  | | ⌧ |  |  |  | | | ⌧ |
| **Tephritidae** |  | ⌧ |  | |  | |  |  |  |  | | |  |
| **Ulidiidae** |  |  |  | |  | |  |  |  |  | | | ⌧ |
| **Xenasteiidae** |  |  |  | |  | |  |  |  |  | | | ⌧ |
| **Xylomyidae** |  |  |  | |  | |  |  |  |  | | | ⌧ |
| **Hymenoptera** | | | | | | | | | | | | | |
| **Aphelinidae** |  |  |  | |  | | ⌧ |  |  |  | | |  |
| **Apidae** |  |  | ⌧ | |  | |  |  |  |  | | |  |
| **Bethylidae** |  |  |  | |  | | ⌧ | ⌧ |  |  | | |  |
| **Braconidae** |  |  |  | |  | | ⌧ |  |  |  | | |  |
| **Ceraphronidae** |  |  |  | |  | | ⌧ |  |  |  | | |  |
| **Chalcidae** |  |  |  | |  | | ⌧ |  |  |  | | |  |
| **Chrysididae** |  |  |  | |  | | ⌧ |  |  |  | | |  |
| **Colletidae** |  |  | ⌧ | |  | |  |  |  |  | | |  |
| **Crabronidae** |  |  |  | |  | |  |  |  |  | | | ⌧ |
| **Diapriidae** |  |  |  | |  | | ⌧ |  |  |  | | |  |
| **Dryinidae** |  |  |  | |  | | ⌧ |  |  |  | | |  |
| **Eulophidae** |  |  |  | |  | | ⌧ |  |  |  | | |  |
| **Eupelmidae** |  |  |  | |  | | ⌧ |  |  |  | | |  |
| **Evaniidae** |  |  |  | |  | | ⌧ |  |  |  | | |  |
| **Figitidae** |  |  |  | |  | | ⌧ |  |  |  | | |  |
| **Formicidae** | *Acropyga* |  |  | |  | |  |  |  |  | | | ⌧ |
| **Formicidae** | *Anochetus* |  |  | |  | |  | ⌧ |  |  | | |  |
| **Formicidae** | *Anoplolepis* |  |  | |  | |  | ⌧ |  |  | | |  |
| **Formicidae** | *Aphaenogaster* |  |  | |  | |  |  |  |  | | | ⌧ |
| **Formicidae** | *Brachyponera* |  |  | |  | |  | ⌧ |  |  | | |  |
| **Formicidae** | *Camponotus* |  |  | |  | |  |  |  |  | | | ⌧ |
| **Formicidae** | *Cardiocondyla* |  |  | |  | |  |  |  |  | | | ⌧ |
| **Formicidae** | *Carebara* |  |  | |  | |  | ⌧ |  |  | | |  |
| **Formicidae** | *Cataulacus* |  |  | |  | |  |  |  |  | | | ⌧ |
| **Formicidae** | *Chronoxenus* |  |  | |  | |  |  |  |  | | | ⌧ |
| **Formicidae** | *Colobopsis* |  |  | |  | |  |  |  |  | | | ⌧ |
| **Formicidae** | *Crematogaster* |  |  | |  | |  |  |  |  | | | ⌧ |
| **Formicidae** | *Cryptopone* |  |  | |  | |  |  |  |  | | | ⌧ |
| **Formicidae** | *Diacamma* |  |  | |  | |  | ⌧ |  |  | | |  |
| **Formicidae** | *Discothyrea* |  |  | |  | |  | ⌧ |  |  | | |  |
| **Formicidae** | *Dolichoderus* |  |  | |  | |  |  |  |  | | | ⌧ |
| **Formicidae** | *Echinopla* |  |  | |  | |  |  |  |  | | | ⌧ |
| **Formicidae** | *Ectomomyrmex* |  |  | |  | |  | ⌧ |  |  | | |  |
| **Formicidae** | *Euponera* |  |  | |  | |  | ⌧ |  |  | | |  |
| **Formicidae** | *Euprenolepis* |  |  | |  | |  |  |  |  | | | ⌧ |
| **Formicidae** | *Gauromyrmex* |  |  | |  | |  |  |  |  | | | ⌧ |
| **Formicidae** | *Hypoponera* |  |  | |  | |  | ⌧ |  |  | | |  |
| **Formicidae** | *Iridomyrmex* |  |  | |  | |  |  |  |  | | | ⌧ |
| **Formicidae** | *Leptogenys* |  |  | |  | |  |  |  |  | | | ⌧ |
| **Formicidae** | *Lioponera* |  |  | |  | |  | ⌧ |  |  | | |  |
| **Formicidae** | *Mayriella* |  |  | |  | |  |  |  |  | | | ⌧ |
| **Formicidae** | *Meranoplus* |  |  | |  | |  |  |  |  | | | ⌧ |
| **Formicidae** | *Mesoponera* |  |  | |  | |  |  |  |  | | | ⌧ |
| **Formicidae** | *Monomorium* |  |  | |  | |  |  |  |  | | | ⌧ |
| **Formicidae** | *Myrmecina* |  |  | |  | |  | ⌧ |  |  | | |  |
| **Formicidae** | *Nylanderia* |  |  | |  | |  |  |  |  | | | ⌧ |
| **Formicidae** | *Odontomachus* |  |  | |  | |  |  |  |  | | | ⌧ |
| **Formicidae** | *Odontoponera* |  |  | |  | |  |  |  |  | | | ⌧ |
| **Formicidae** | *Oecophylla* |  |  | |  | |  |  |  |  | | | ⌧ |
| **Formicidae** | *Paraparatrechina* |  |  | |  | |  |  |  |  | | | ⌧ |
| **Formicidae** | *Paratopula* |  |  | |  | |  |  |  |  | | | ⌧ |
| **Formicidae** | *Paratrechina* |  |  | |  | |  |  |  |  | | | ⌧ |
| **Formicidae** | *Pheidole* |  |  | |  | |  |  |  |  | | | ⌧ |
| **Formicidae** | *Philidris* | ⌧ |  | |  | |  |  |  |  | | |  |
| **Formicidae** | *Platythyrea* |  |  | |  | |  | ⌧ |  |  | | |  |
| **Formicidae** | *Polyrhachis* | ⌧ |  | |  | |  |  |  |  | | |  |
| **Formicidae** | *Ponera* |  |  | |  | |  |  |  |  | | | ⌧ |
| **Formicidae** | *Prenolepis* |  |  | |  | |  |  |  |  | | | ⌧ |
| **Formicidae** | *Prionopelta* |  |  | |  | |  |  |  |  | | | ⌧ |
| **Formicidae** | *Proatta* |  |  | |  | |  | ⌧ |  |  | | |  |
| **Formicidae** | *Probolomyrmex* |  |  | |  | |  |  |  |  | | | ⌧ |
| **Formicidae** | *Pseudoneoponera* |  |  | |  | |  | ⌧ |  |  | | |  |
| **Formicidae** | *Strumigenys* |  |  | |  | |  | ⌧ |  |  | | |  |
| **Formicidae** | *Rhopalomastix* |  |  | |  | |  | ⌧ |  |  | | |  |
| **Formicidae** | *Solenopsis* |  |  | |  | |  |  |  |  | | | ⌧ |
| **Formicidae** | *Stigmatomma* |  |  | |  | |  | ⌧ |  |  | | |  |
| **Formicidae** | *Strumigenys* |  |  | |  | |  | ⌧ |  |  | | |  |
| **Formicidae** | *Tapinoma* |  |  | |  | |  |  |  |  | | | ⌧ |
| **Formicidae** | *Technomyrmex* |  |  | |  | |  |  |  |  | | | ⌧ |
| **Formicidae** | *Tetramorium* |  |  | |  | |  | ⌧ |  |  | | |  |
| **Formicidae** | *Tetraponera* | ⌧ |  | |  | |  |  |  |  | | |  |
| **Formicidae** | *Vollenhovia* |  |  | |  | |  |  |  |  | | | ⌧ |
| **Halictidae** |  |  | ⌧ | |  | |  |  |  |  | | |  |
| **Ichneumonidae** |  |  |  | |  | | ⌧ |  |  |  | | |  |
| **Megachilidae** |  |  | ⌧ | |  | |  |  |  |  | | |  |
| **Mymaridae** |  |  |  | |  | | ⌧ |  |  |  | | |  |
| **Platygastridae** |  |  |  | |  | | ⌧ |  |  |  | | |  |
| **Pompilidae** |  |  |  | |  | | ⌧ |  |  |  | | |  |
| **Pteromalidae** |  |  |  | |  | | ⌧ |  |  |  | | |  |
| **Scoliidae** |  |  |  | |  | | ⌧ |  |  |  | | |  |
| **Sphecidae** |  |  |  | |  | | ⌧ |  |  |  | | |  |
| **Sphecidae** |  |  |  | |  | |  | ⌧ |  |  | | |  |
| **Tiphiidae** |  |  |  | |  | | ⌧ |  |  |  | | |  |
| **Trichogrammatidae** |  |  |  | |  | | ⌧ |  |  |  | | |  |
| **Vespidae** |  |  | ⌧ | |  | |  | ⌧ |  |  | | |  |

**Table S4.** Number of specimens from Singapore, Hong Kong and Brunei, as well as the size of the randomized subsample from Singapore.

|  | No. of Specimens | | | |  |
| --- | --- | --- | --- | --- | --- |
| Taxon | **Singapore** | **Singapore (Rarefied)** | **Hong Kong** | **Brunei** | **Thailand** |
| Dolichopodidae | 17860 | 2800 | 2563 | 2798 | 924 |
| Phoridae | 2134 | 560 | 562 | 272 | - |
| Mycetophilidae | 223 | 180 | 186 | - | - |
| Total | **20217** | **3540** | **3311** | **3070** | **924** |

**Table S5.** Number of species of vascular plants for each sampling site in Singapore from checklist data.

| Sampling Site | Habitat | No. of Plant Species | Reference |
| --- | --- | --- | --- |
| Nee Soon freshwater swamp | Freshwater swamp forest | 1150 | Wong et al., 2013[81] |
| Bukit Timah Nature Reserve | Rainforest | 1250 | Ho et al., 2019[69] |
| Kent Ridge | Urban-edge/disturbed forest | 420 | Tan et al., 2019[116] |
| Pulau Ubin | Mangrove | 245 | Lee et al., 2003[75] |
| Sungei Buloh Wetland Reserve | Mangrove | 249 | Tan et al., 1997[76] |
| Pulau Semakau | Mangrove | 165 | Teo et al., 2011[77] |

**Table S6.** Number and distribution of mOTUs delimited using different thresholds (144,865 barcoded specimens)

| Habitat/Country | No. of Barcodes | No. of mOTUs from Objective Clustering | | | No. of mOTUs from USEARCH | | |
| --- | --- | --- | --- | --- | --- | --- | --- |
|  |  | **2%** | **3%** | **4%** | **id=0.98** | **id=0.97** | **id=0.96** |
| Singapore full dataset | | | | | | | |
| Mangroves | 67239 | 3557 | 3437 | 3320 | 3710 | 3524 | 3436 |
| Rainforest | 15669 | 2625 | 2573 | 2539 | 2669 | 2603 | 2570 |
| Swamp forest | 9464 | 1843 | 1804 | 1753 | 1895 | 1828 | 1795 |
| Urban forest | 20323 | 1552 | 1515 | 1478 | 1616 | 1549 | 1510 |
| Freshwater swamp | 21994 | 1881 | 1812 | 1744 | 1988 | 1878 | 1805 |
| Coastal forest | 9118 | 1707 | 1667 | 1627 | 1755 | 1691 | 1664 |
| Total | **143807** | **8903** | **8572** | **8256** | **9315** | **8821** | **8520** |
| Subset used for guild-level analysis | | | | | | | |
| Mangroves | 37641 | 1778 | 1720 | 1673 | 1828 | 1744 | 1702 |
| Rainforest | 9212 | 1525 | 1490 | 1474 | 1545 | 1503 | 1483 |
| Swamp forest | 5893 | 1090 | 1052 | 1030 | 1105 | 1070 | 1048 |
| Urban forest | 9320 | 919 | 898 | 885 | 941 | 908 | 893 |
| Total | **62066** | **4169** | **4002** | **3917** | **4298** | **4098** | **3994** |
| Southeast and East Asian datasets | | | | | | | |
| *Dolichopodidae* | | | | | | | |
| Singapore | 17860 | 263 | 254 | 248 | 280 | 259 | 249 |
| Hong Kong | 2601 | 111 | 109 | 104 | 115 | 110 | 106 |
| Brunei | 2800 | 98 | 96 | 95 | 107 | 98 | 95 |
| Thailand | 924 | 80 | 74 | 72 | 93 | 80 | 73 |
| Total | **24185** | **480** | **453** | **426** | **543** | **482** | **447** |
| *Phoridae* | | | | | | | |
| Singapore | 2134 | 293 | 281 | 278 | 300 | 285 | 280 |
| Hong Kong | 562 | 137 | 129 | 125 | 138 | 130 | 129 |
| Brunei | 272 | 76 | 76 | 75 | 77 | 76 | 75 |
| Total | **2968** | **453** | **429** | **417** | **467** | **437** | **431** |
| *Mycetophilidae* | | | | | | | |
| Singapore | 223 | 45 | 44 | 43 | 45 | 44 | 44 |
| Hong Kong | 186 | 26 | 25 | 25 | 26 | 25 | 25 |
| Total | **409** | **69** | **67** | **67** | **70** | **67** | **67** |

**Table S7.** Common and rare species found in only 1, 2, 3, 4, 5 or all habitats.

|  | No. of species | | | | |
| --- | --- | --- | --- | --- | --- |
|  | **Full dataset** | **No singletons** | **No doubletons** | **No species with <5 specimens** | **No species with <10 specimens** |
| Species in mangroves only | 1788 | 880 | 638 | 441 | 256 |
| Species in rainforests only | 1569 | 638 | 415 | 243 | 91 |
| Species in swamp forests only | 875 | 342 | 200 | 102 | 39 |
| Species in urban forests only | 509 | 166 | 101 | 58 | 25 |
| Species in freshwater swamps only | 794 | 360 | 237 | 127 | 56 |
| Species in coastal forests only | 454 | 153 | 71 | 33 | 14 |
| Species in two habitats | 1580 | 1580 | 1253 | 887 | 555 |
| Species in three habitats | 565 | 565 | 565 | 494 | 350 |
| Species in four habitats | 274 | 274 | 274 | 265 | 230 |
| Species in five habitats | 116 | 116 | 116 | 116 | 109 |
| Species in all habitats | 48 | 48 | 48 | 48 | 48 |
| Total | **8572** | **5122** | **3918** | **2814** | **1773** |

**Table S8.** Species turnover ANOSIM analysis results indicate distinct communities in each habitat type, whether with singletons and doubletons removed, or species with less than 5 and 10 specimens. Pairwise p-value outputs are displayed in the bottom-left of the pairwise matrix while the R-statistics are displayed at the top-right.

**No Singletons**

| **Overall P:** 0.001 **Overall R:** 0.777 | | | | |  |  |
| --- | --- | --- | --- | --- | --- | --- |
|  | **Rainforest** | **Urban forest** | **Swamp forest** | **Mangrove** | **Freshwater swamp** | **Coastal forest** |
| **Rainforest** |  | 0.809 | 0.981 | 0.948 | 0.972 | 0.951 |
| **Urban forest** | 0.001 |  | 0.747 | 0.815 | 0.571 | 0.173 |
| **Swamp forest** | 0.001 | 0.001 |  | 0.927 | 0.756 | 0.893 |
| **Mangrove** | 0.001 | 0.001 | 0.001 |  | 0.852 | 0.541 |
| **Freshwater swamp** | 0.001 | 0.001 | 0.008 | 0.001 |  | 0.347 |
| **Coastal forest** | 0.001 | 0.083 | 0.005 | 0.001 | 0.017 |  |

**No Doubletons**

| **Overall P:** 0.001 **Overall R:** 0.774 | | | | |  |  |
| --- | --- | --- | --- | --- | --- | --- |
|  | **Rainforest** | **Urban forest** | **Swamp forest** | **Mangrove** | **Freshwater swamp** | **Coastal forest** |
| **Rainforest** |  | 0.803 | 0.980 | 0.946 | 0.972 | 0.954 |
| **Urban forest** | 0.001 |  | 0.735 | 0.816 | 0.563 | 0.179 |
| **Swamp forest** | 0.001 | 0.001 |  | 0.922 | 0.750 | 0.889 |
| **Mangrove** | 0.001 | 0.001 | 0.001 |  | 0.849 | 0.538 |
| **Freshwater swamp** | 0.001 | 0.001 | 0.008 | 0.001 |  | 0.331 |
| **Coastal forest** | 0.002 | 0.072 | 0.005 | 0.001 | 0.019 |  |

**No Species <5 Specimens**

| **Overall P:** 0.001 **Overall R:** 0.767 | | | | |  |  |
| --- | --- | --- | --- | --- | --- | --- |
|  | **Rainforest** | **Urban forest** | **Swamp forest** | **Mangrove** | **Freshwater swamp** | **Coastal forest** |
| **Rainforest** |  | 0.795 | 0.970 | 0.941 | 0.971 | 0.954 |
| **Urban forest** | 0.001 |  | 0.720 | 0.817 | 0.559 | 0.180 |
| **Swamp forest** | 0.001 | 0.001 |  | 0.913 | 0.750 | 0.885 |
| **Mangrove** | 0.001 | 0.001 | 0.001 |  | 0.843 | 0.533 |
| **Freshwater swamp** | 0.002 | 0.001 | 0.008 | 0.001 |  | 0.331 |
| **Coastal forest** | 0.002 | 0.061 | 0.005 | 0.001 | 0.017 |  |

**No Species <10 Specimens**

| **Overall P:** 0.001 **Overall R:** 0.759 | | | | |  |  |
| --- | --- | --- | --- | --- | --- | --- |
|  | **Rainforest** | **Urban forest** | **Swamp forest** | **Mangrove** | **Freshwater swamp** | **Coastal forest** |
| **Rainforest** |  | 0.779 | 0.959 | 0.934 | 0.967 | 0.952 |
| **Urban forest** | 0.001 |  | 0.701 | 0.819 | 0.548 | 0.178 |
| **Swamp forest** | 0.001 | 0.002 |  | 0.904 | 0.738 | 0.877 |
| **Mangrove** | 0.001 | 0.001 | 0.001 |  | 0.837 | 0.526 |
| **Freshwater swamp** | 0.002 | 0.001 | 0.008 | 0.001 |  | 0.331 |
| **Coastal forest** | 0.001 | 0.062 | 0.005 | 0.001 | 0.017 |  |

**Table S9.** Species turnover SIMPER analysis results indicate distinct communities in each habitat type, whether with singletons and doubletons removed, or species with less than 5 and 10 specimens.

**No Singletons**

|  | **Within habitat (%)** | **Between habitats (%)** | | | | | |
| --- | --- | --- | --- | --- | --- | --- | --- |
|  |  | **Rain- forest** | **Urban forest** | **Swamp forest** | **Mangrove** | **Fresh-water swamp** | **Coastal forest** |
| **Rainforest** | 33.65 |  |  |  |  |  |  |
| **Urban forest** | 13.70 | 3.57 |  |  |  |  |  |
| **Swamp forest** | 35.74 | 15.86 | 3.31 |  |  |  |  |
| **Mangrove** | 12.78 | 1.80 | 3.26 | 2.22 |  |  |  |
| **Freshwater swamp** | 18.80 | 2.36 | 5.06 | 4.57 | 2.93 |  |  |
| **Coastal forest** | 12.98 | 4.27 | 10.04 | 4.50 | 6.44 | 9.82 |  |

**No Doubletons**

|  | **Within habitat (%)** | **Between habitats (%)** | | | | | |
| --- | --- | --- | --- | --- | --- | --- | --- |
|  |  | **Rain- forest** | **Urban forest** | **Swamp forest** | **Mangrove** | **Fresh-water swamp** | **Coastal forest** |
| **Rainforest** | 35.83 |  |  |  |  |  |  |
| **Urban forest** | 14.15 | 3.79 |  |  |  |  |  |
| **Swamp forest** | 38.05 | 17.12 | 3.55 |  |  |  |  |
| **Mangrove** | 13.14 | 1.91 | 3.36 | 2.39 |  |  |  |
| **Freshwater swamp** | 19.61 | 2.52 | 5.30 | 4.90 | 3.07 |  |  |
| **Coastal forest** | 13.62 | 4.57 | 10.44 | 4.86 | 6.68 | 10.37 |  |

**No Species <5 Specimens**

|  | **Within habitat (%)** | **Between habitats (%)** | | | | | |
| --- | --- | --- | --- | --- | --- | --- | --- |
|  |  | **Rain- forest** | **Urban forest** | **Swamp forest** | **Mangrove** | **Fresh-water swamp** | **Coastal forest** |
| **Rainforest** | 38.68 |  |  |  |  |  |  |
| **Urban forest** | 14.86 | 4.13 |  |  |  |  |  |
| **Swamp forest** | 40.06 | 18.89 | 3.93 |  |  |  |  |
| **Mangrove** | 13.65 | 2.08 | 3.50 | 2.65 |  |  |  |
| **Freshwater swamp** | 20.84 | 2.76 | 5.68 | 5.46 | 3.29 |  |  |
| **Coastal forest** | 14.49 | 4.99 | 11.08 | 5.39 | 7.03 | 11.15 |  |

**No Species <10 Specimens**

|  | **Within habitat (%)** | **Between habitats (%)** | | | | | |
| --- | --- | --- | --- | --- | --- | --- | --- |
|  |  | **Rain- forest** | **Urban forest** | **Swamp forest** | **Mangrove** | **Fresh-water swamp** | **Coastal forest** |
| **Rainforest** | 42.79 |  |  |  |  |  |  |
| **Urban forest** | 15.91 | 4.79 |  |  |  |  |  |
| **Swamp forest** | 42.93 | 21.56 | 4.49 |  |  |  |  |
| **Mangrove** | 14.55 | 2.41 | 3.75 | 3.04 |  |  |  |
| **Freshwater swamp** | 22.41 | 3.25 | 6.28 | 6.25 | 3.63 |  |  |
| **Coastal forest** | 15.95 | 5.83 | 12.09 | 6.19 | 7.65 | 12.16 |  |

**Table S10.** Species turnover and nestedness analysis reveal that the high dissimilarity is due more to turnover rather than nestedness, whether with singletons and doubletons removed, or species with less than 5 and 10 specimens. Pairwise turnover values are displayed in the bottom-left of the pairwise matrix while the nestedness values are in the top-right.

**No Singletons**

| **Overall Dissimilarity:** 0.944 **Overall** **Turnover:** 0.894 **Overall Nestedness:** 0.051 | | | | | | |
| --- | --- | --- | --- | --- | --- | --- |
|  | **Rainforest** | **Urban forest** | **Swamp forest** | **Mangrove** | **Freshwater swamp** | **Coastal forest** |
| **Rainforest** |  | 0.013 | 0.075 | 0.058 | 0.009 | 0.020 |
| **Urban forest** | 0.911 |  | 0.031 | 0.099 | 0.005 | 0.107 |
| **Swamp forest** | 0.693 | 0.918 |  | 0.098 | 0.030 | 0.002 |
| **Mangrove** | 0.908 | 0.816 | 0.871 |  | 0.063 | 0.263 |
| **Freshwater swamp** | 0.953 | 0.889 | 0.928 | 0.876 |  | 0.098 |
| **Coastal forest** | 0.905 | 0.695 | 0.936 | 0.648 | 0.748 |  |

**No Doubletons**

| **Overall Dissimilarity:** 0.944 **Overall** **Turnover:** 0.892 **Overall Nestedness:** 0.052 | | | | | | |
| --- | --- | --- | --- | --- | --- | --- |
|  | **Rainforest** | **Urban forest** | **Swamp forest** | **Mangrove** | **Freshwater swamp** | **Coastal forest** |
| **Rainforest** |  | 0.015 | 0.078 | 0.060 | 0.010 | 0.020 |
| **Urban forest** | 0.909 |  | 0.033 | 0.099 | 0.004 | 0.112 |
| **Swamp forest** | 0.685 | 0.915 |  | 0.102 | 0.032 | 0.003 |
| **Mangrove** | 0.906 | 0.816 | 0.868 |  | 0.064 | 0.268 |
| **Freshwater swamp** | 0.952 | 0.889 | 0.926 | 0.875 |  | 0.101 |
| **Coastal forest** | 0.903 | 0.687 | 0.934 | 0.643 | 0.744 |  |

**No Species <5 Specimens**

| **Overall Dissimilarity:** 0.944 **Overall** **Turnover:** 0.891 **Overall Nestedness:** 0.054 | | | | | | |
| --- | --- | --- | --- | --- | --- | --- |
|  | **Rainforest** | **Urban forest** | **Swamp forest** | **Mangrove** | **Freshwater swamp** | **Coastal forest** |
| **Rainforest** |  | 0.017 | 0.081 | 0.063 | 0.011 | 0.020 |
| **Urban forest** | 0.905 |  | 0.037 | 0.099 | 0.004 | 0.118 |
| **Swamp forest** | 0.677 | 0.912 |  | 0.107 | 0.035 | 0.004 |
| **Mangrove** | 0.904 | 0.817 | 0.862 |  | 0.064 | 0.274 |
| **Freshwater swamp** | 0.950 | 0.889 | 0.922 | 0.875 |  | 0.103 |
| **Coastal forest** | 0.902 | 0.679 | 0.931 | 0.638 | 0.741 |  |

**No Species <10 Specimens**

| **Overall Dissimilarity:** 0.945 **Overall** **Turnover:** 0.888 **Overall Nestedness:** 0.057 | | | | | | |
| --- | --- | --- | --- | --- | --- | --- |
|  | **Rainforest** | **Urban forest** | **Swamp forest** | **Mangrove** | **Freshwater swamp** | **Coastal forest** |
| **Rainforest** |  | 0.023 | 0.082 | 0.069 | 0.014 | 0.018 |
| **Urban forest** | 0.897 |  | 0.041 | 0.099 | 0.003 | 0.128 |
| **Swamp forest** | 0.665 | 0.907 |  | 0.115 | 0.038 | 0.005 |
| **Mangrove** | 0.898 | 0.818 | 0.856 |  | 0.065 | 0.282 |
| **Freshwater swamp** | 0.944 | 0.890 | 0.917 | 0.876 |  | 0.105 |
| **Coastal forest** | 0.899 | 0.665 | 0.927 | 0.632 | 0.742 |  |

**Table S11.** Number of species from each guild, site and habitat type.

| Guild | Rainforest | Swamp Forest | Urban Forest | Mangrove Forest | | | |
| --- | --- | --- | --- | --- | --- | --- | --- |
|  | **Bukit Timah** | **Nee Soon** | **Kent Ridge** | **Pulau Ubin** | **Sungei Buloh** | **Pulau Semakau (Old)** | **Pulau Semakau (New)** |
| Phytophages | 471 | 447 | 125 | 155 | 110 | 109 | 71 |
| Pollinators | 15 | 17 | 1 | 29 | 29 | 20 | 20 |
| Fungivores | 400 | 380 | 83 | 67 | 46 | 27 | 12 |
| Parasitoids | 164 | 78 | 133 | 83 | 71 | 61 | 38 |
| Predators | 153 | 135 | 83 | 219 | 136 | 167 | 117 |
| Haematophages | 33 | 49 | 0 | 65 | 57 | 40 | 24 |
| Detritivores | 74 | 68 | 17 | 90 | 81 | 58 | 43 |

**Table S12.** Species turnover ANOSIM analysis results indicate distinct communities in each habitat type for each ecological guild. Pairwise p-value outputs are displayed in the bottom-left of the pairwise matrix while the R-statistics are displayed at the top-right.

**Phytophages**

| **Overall P:** 0.001 **Overall R:** 0.588 | | | | |
| --- | --- | --- | --- | --- |
|  | **Rainforest** | **Urban forest** | **Swamp forest** | **Mangrove** |
| **Rainforest** |  | 1.000 | 0.706 | 0.721 |
| **Urban forest** | 0.001 |  | 1.000 | 0.468 |
| **Swamp forest** | 0.001 | 0.029 |  | 0.665 |
| **Mangrove** | 0.001 | 0.003 | 0.001 |  |

**Pollinators**

| **Overall P:** 0.001 **Overall R:** 0.836 | | | |
| --- | --- | --- | --- |
|  | **Rainforest** | **Swamp forest** | **Mangrove** |
| **Rainforest** |  | 0.387 | 0.915 |
| **Swamp forest** | 0.127 |  | 0.853 |
| **Mangrove** | 0.001 | 0.004 |  |

**Fungivores**

| **Overall P:** 0.001 **Overall R:** 0.351 | | | | |
| --- | --- | --- | --- | --- |
|  | **Rainforest** | **Urban forest** | **Swamp forest** | **Mangrove** |
| **Rainforest** |  | 1.000 | 0.726 | 0.435 |
| **Urban forest** | 0.001 |  | 1.000 | 0.088 |
| **Swamp forest** | 0.001 | 0.029 |  | 0.432 |
| **Mangrove** | 0.001 | 0.206 | 0.001 |  |

**Parasitoids**

| **Overall P:** 0.001 **Overall R:** 0.758 | | | | |
| --- | --- | --- | --- | --- |
|  | **Rainforest** | **Urban forest** | **Swamp forest** | **Mangrove** |
| **Rainforest** |  | 0.962 | 0.925 | 0.793 |
| **Urban forest** | 0.001 |  | 1.000 | 0.736 |
| **Swamp forest** | 0.018 | 0.067 |  | 0.711 |
| **Mangrove** | 0.001 | 0.001 | 0.006 |  |

**Predators**

| **Overall P:** 0.001 **Overall R:** 0.906 | | | | |
| --- | --- | --- | --- | --- |
|  | **Rainforest** | **Urban forest** | **Swamp forest** | **Mangrove** |
| **Rainforest** |  | 1.000 | 0.414 | 0.954 |
| **Urban forest** | 0.001 |  | 1.000 | 0.916 |
| **Swamp forest** | 0.109 | 0.067 |  | 0.913 |
| **Mangrove** | 0.001 | 0.001 | 0.002 |  |

**Haematophages**

| **Overall P:** 0.001 **Overall R:** 0.905 | | | |
| --- | --- | --- | --- |
|  | **Rainforest** | **Swamp forest** | **Mangrove** |
| **Rainforest** |  | 0.435 | 0.957 |
| **Swamp forest** | 0.139 |  | 0.791 |
| **Mangrove** | 0.001 | 0.002 |  |

**Detritivores**

| **Overall P:** 0.001 **Overall R:** 0.853 | | | | |
| --- | --- | --- | --- | --- |
|  | **Rainforest** | **Urban forest** | **Swamp forest** | **Mangrove** |
| **Rainforest** |  | 0.613 | 0.487 | 0.949 |
| **Urban forest** | 0.008 |  | 1.000 | 0.904 |
| **Swamp forest** | 0.056 | 0.100 |  | 0.614 |
| **Mangrove** | 0.001 | 0.001 | 0.002 |  |

**Table S13.** Species turnover SIMPER analysis results indicate distinct communities in each habitat type for each ecological guild.

**Phytophages**

|  | **Within habitat (%)** | **Between habitats (%)** | | | |
| --- | --- | --- | --- | --- | --- |
|  |  | **Rainforest** | **Urban forest** | **Swamp forest** | **Mangrove** |
| **Rainforest** | 30.46 |  |  |  |  |
| **Urban forest** | 27.72 | 4.93 |  |  |  |
| **Swamp forest** | 34.21 | 18.38 | 5.18 |  |  |
| **Mangrove** | 12.37 | 1.29 | 4.13 | 1.67 |  |

**Pollinators**

|  | **Within habitat (%)** | **Between habitats (%)** | | |
| --- | --- | --- | --- | --- |
|  |  | **Rainforest** | **Swamp forest** | **Mangrove** |
| **Rainforest** | 41.27 |  |  |  |
| **Swamp forest** | 48.30 | 28.15 |  |  |
| **Mangrove** | 26.01 | 0.88 | 3.57 |  |

**Fungivores**

|  | **Within habitat (%)** | **Between habitats (%)** | | | |
| --- | --- | --- | --- | --- | --- |
|  |  | **Rainforest** | **Urban forest** | **Swamp forest** | **Mangrove** |
| **Rainforest** | 32.01 |  |  |  |  |
| **Urban forest** | 31.71 | 4.48 |  |  |  |
| **Swamp forest** | 36.88 | 19.40 | 4.61 |  |  |
| **Mangrove** | 10.58 | 1.87 | 8.26 | 1.34 |  |

**Parasitoids**

|  | **Within habitat (%)** | **Between habitats (%)** | | | |
| --- | --- | --- | --- | --- | --- |
|  |  | **Rainforest** | **Urban forest** | **Swamp forest** | **Mangrove** |
| **Rainforest** | 27.47 |  |  |  |  |
| **Urban forest** | 10.13 | 3.18 |  |  |  |
| **Swamp forest** | 59.26 | 10.76 | 1.14 |  |  |
| **Mangrove** | 12.00 | 2.40 | 2.43 | 2.84 |  |

**Predators**

|  | **Within habitat (%)** | **Between habitats (%)** | | | |
| --- | --- | --- | --- | --- | --- |
|  |  | **Rainforest** | **Urban forest** | **Swamp forest** | **Mangrove** |
| **Rainforest** | 29.22 |  |  |  |  |
| **Urban forest** | 34.03 | 4.60 |  |  |  |
| **Swamp forest** | 64.88 | 19.72 | 3.62 |  |  |
| **Mangrove** | 22.78 | 0.28 | 1.35 | 1.20 |  |

**Haematophages**

|  | **Within habitat (%)** | **Between habitats (%)** | | |
| --- | --- | --- | --- | --- |
|  |  | **Rainforest** | **Swamp forest** | **Mangrove** |
| **Rainforest** | 18.86 |  |  |  |
| **Swamp forest** | 56.42 | 10.55 |  |  |
| **Mangrove** | 27.40 | 0.61 | 9.27 |  |

**Detritivores**

|  | **Within habitat (%)** | **Between habitats (%)** | | | |
| --- | --- | --- | --- | --- | --- |
|  |  | **Rainforest** | **Urban forest** | **Swamp forest** | **Mangrove** |
| **Rainforest** | 14.99 |  |  |  |  |
| **Urban forest** | 20.18 | 7.12 |  |  |  |
| **Swamp forest** | 52.77 | 9.49 | 1.67 |  |  |
| **Mangrove** | 18.87 | 0.37 | 1.33 | 6.91 |  |

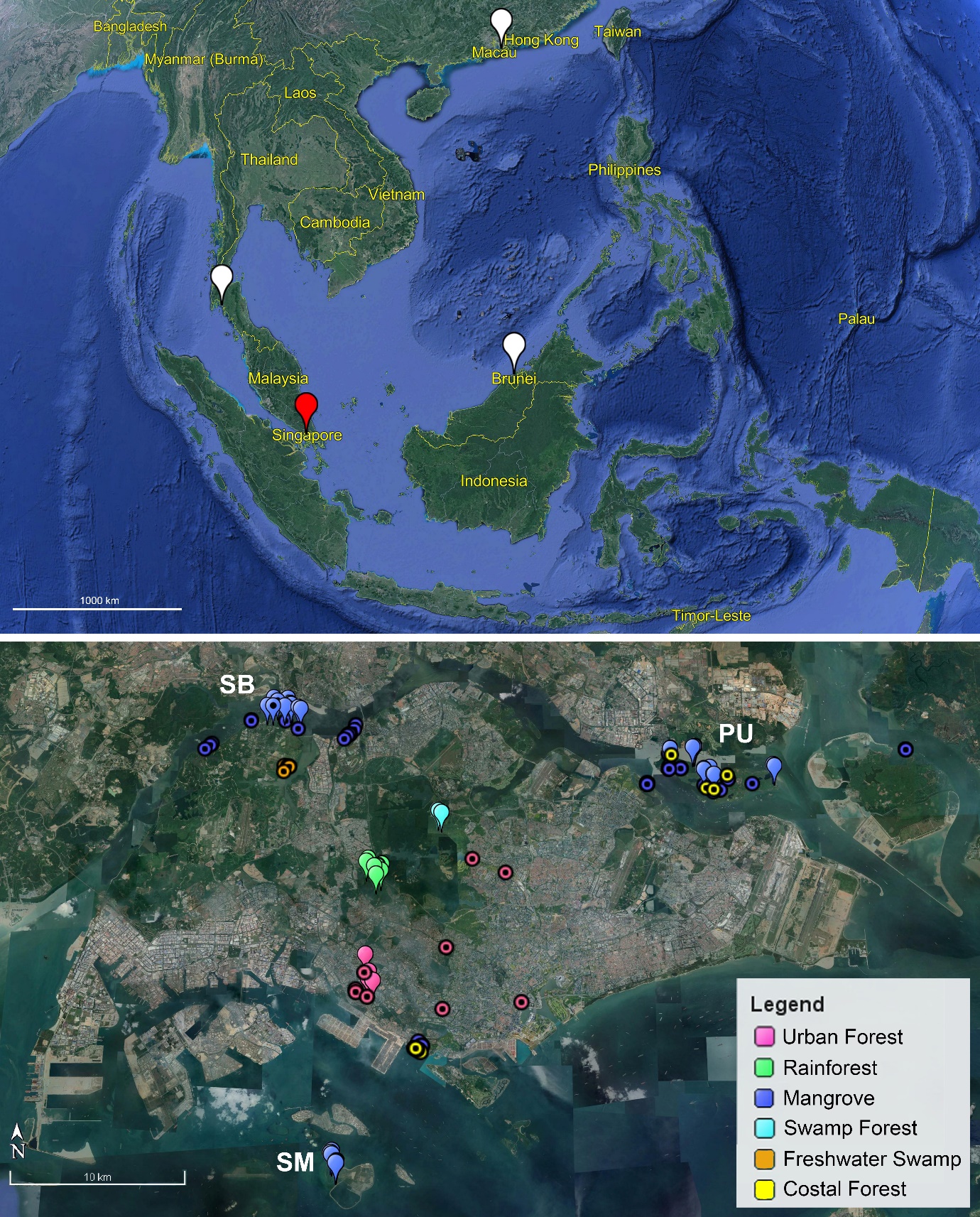

**Figure S1.** Sampling localities in the Oriental Realm (*top*: Singapore, red; other countries, white) and within Singapore (*bottom*: circular markers indicate trapping sites excluded from the species turnover analyses; pin markers with dot indicate traps excluded from guild-level analyses).

**Figure S2.** Arthropod orders sampled with Malaise traps in this study and their species proportions. The number beside each order indicates the number of species sampled based on 3% p-distance objective clustering.

|  |  |
| --- | --- |
| Diptera (5226) | 60.9659% |
| Hymenoptera (1152) | 13.4391% |
| Hemiptera (733) | 8.5511% |
| Lepidoptera (463) | 5.40131% |
| Araneae (374) | 4.36304% |
| Coleoptera (315) | 3.67476% |
| Blattodea (141) | 1.64489% |
| Psocodea (69) | 0.804946% |
| Orthoptera (61) | 0.711619% |
| Neuroptera (14) | 0.163322% |
| Mantodea (13) | 0.151657% |
| Trichoptera (10) | 0.116659% |
| Dermaptera (1) | 0.0116659% |

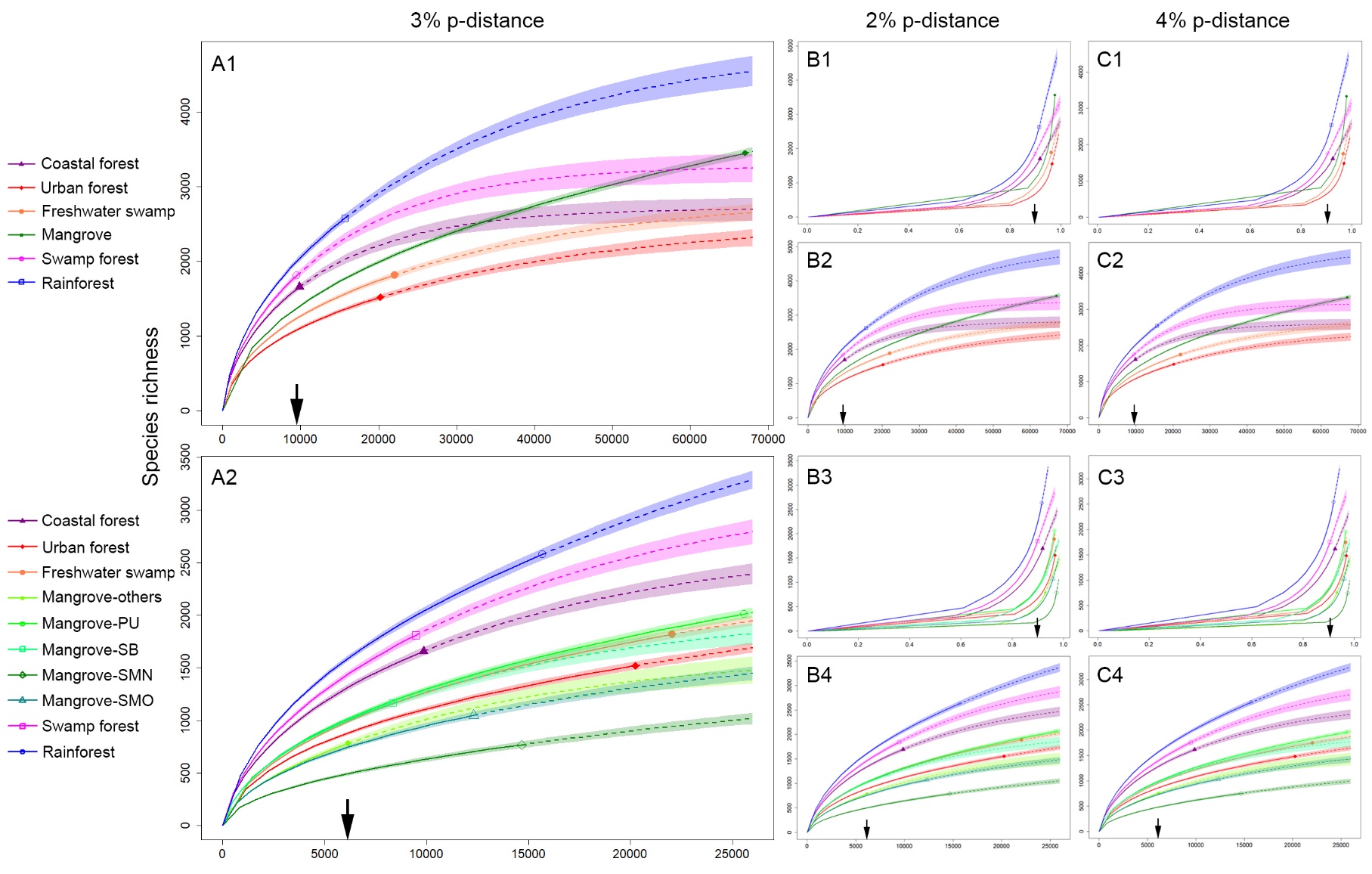

**Figure S3.** Insect alpha-diversity across tropical forest habitats rarefied by specimens (A1 & 2, B2 & 4, C2 & 4) and coverage (B1 & 3, C1 & 3), for 2% (B1 – 4), 3% (A1 – 2) and 4% (C1 – 4) p-distances mOTUs. Mangroves are treated as a single habitat (top) and split by site in a separate analysis (bottom): Pulau Ubin (PU), Sungei Buloh (SB), Pulau Semakau old grove (SMO), Pulau Semakau new grove (SMN); solid lines = rarefaction; dotted = extrapolations. The arrow on the x-axis indicate the point of rarefaction at which species richness comparisons were made.

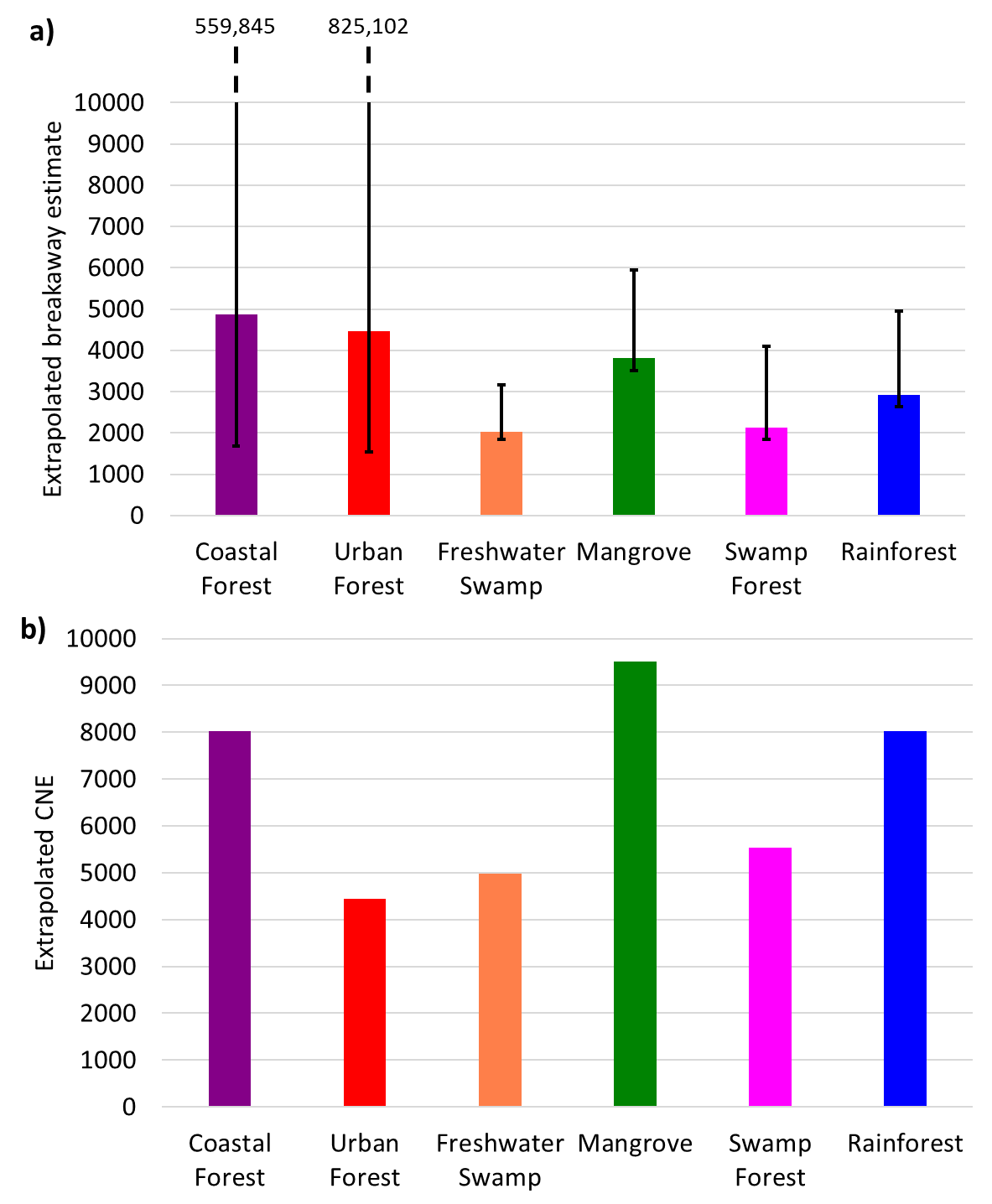

**Figure S4.** Species estimates for each habitat based on a) breakaway estimates and b) CNE extrapolation find mangroves more species-rich than swamp forest and rainforest habitats.

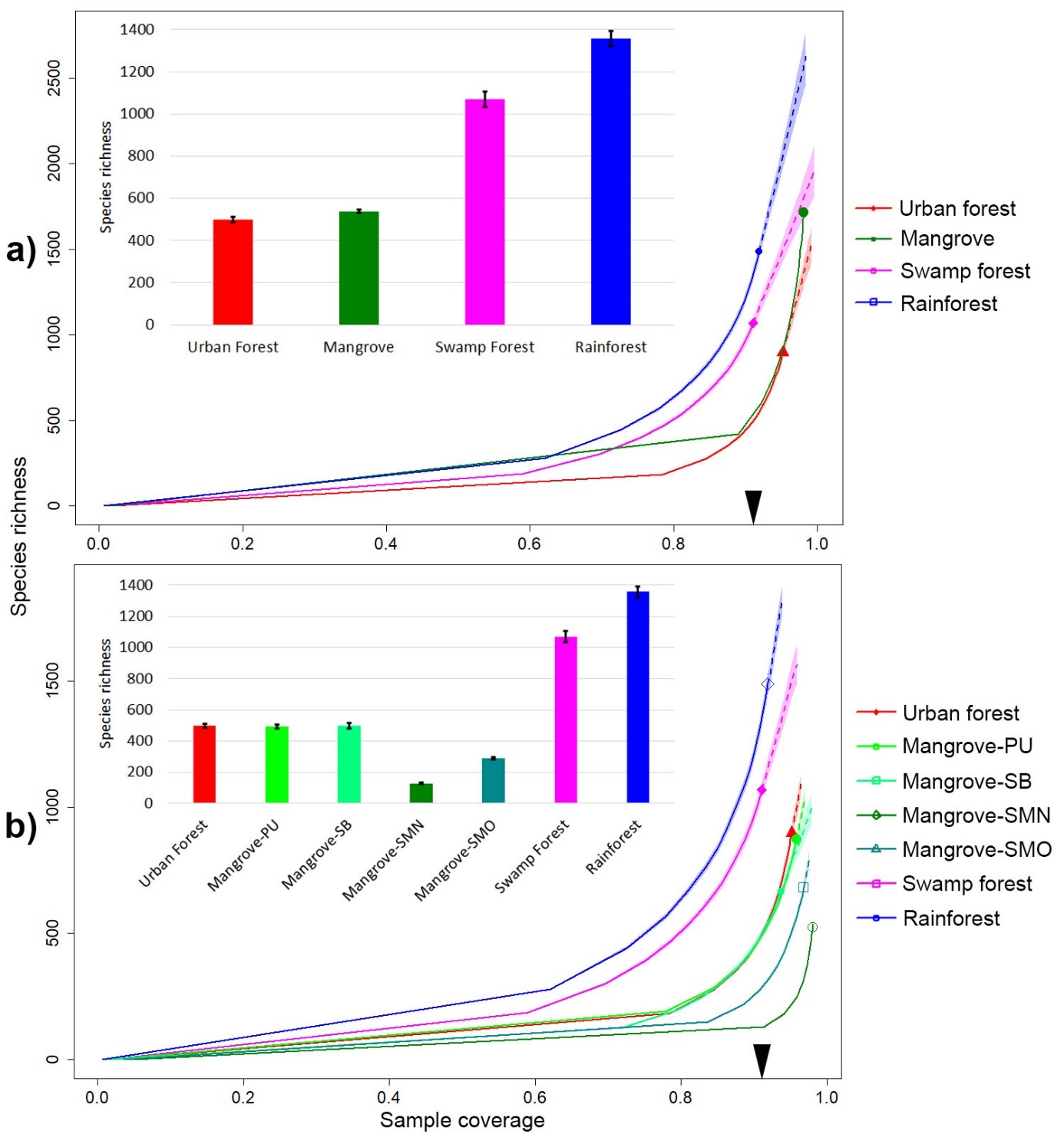

**Figure S5.** Insect alpha-diversity across tropical forest habitats for the core dataset. (a) Mangroves treated as one habitat; (b) Comparison of mangrove sites: Pulau Ubin (PU), Sungei Buloh (SB), Pulau Semakau old-growth (SMO), Pulau Semakau new-growth (SMN); solid lines = rarefaction; dotted = extrapolations. The arrow on the x-axis indicates the point of rarefaction where species richness comparisons were made, which is reflected in the bar charts with associated 95% confidence intervals.

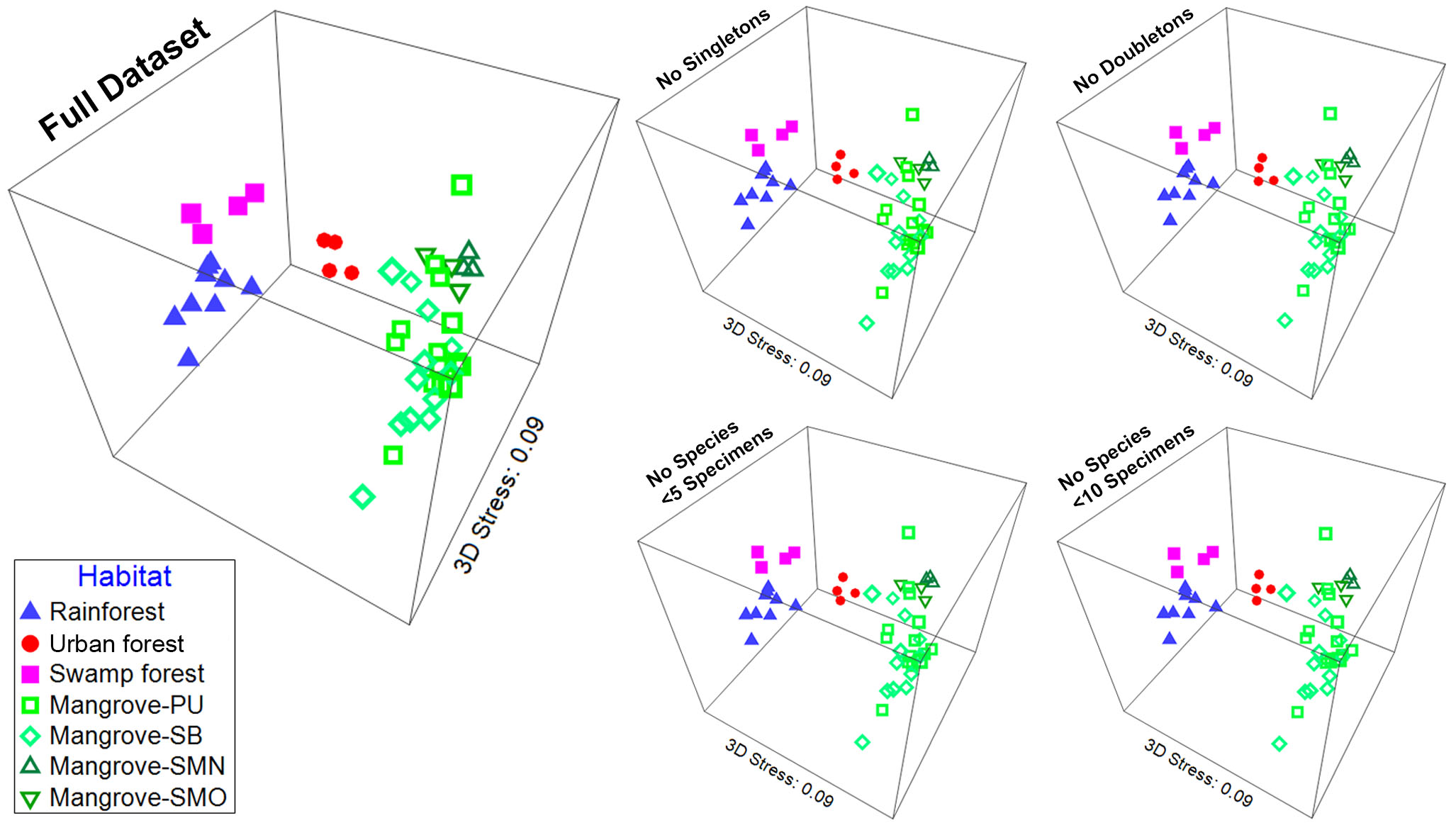

**Figure S6.** Insect communities from the core dataset are distinct across tropical forest habitats based on Bray-Curtis distances. as illustrated with 3D NMDS plots, regardless of whether rare species are removed.

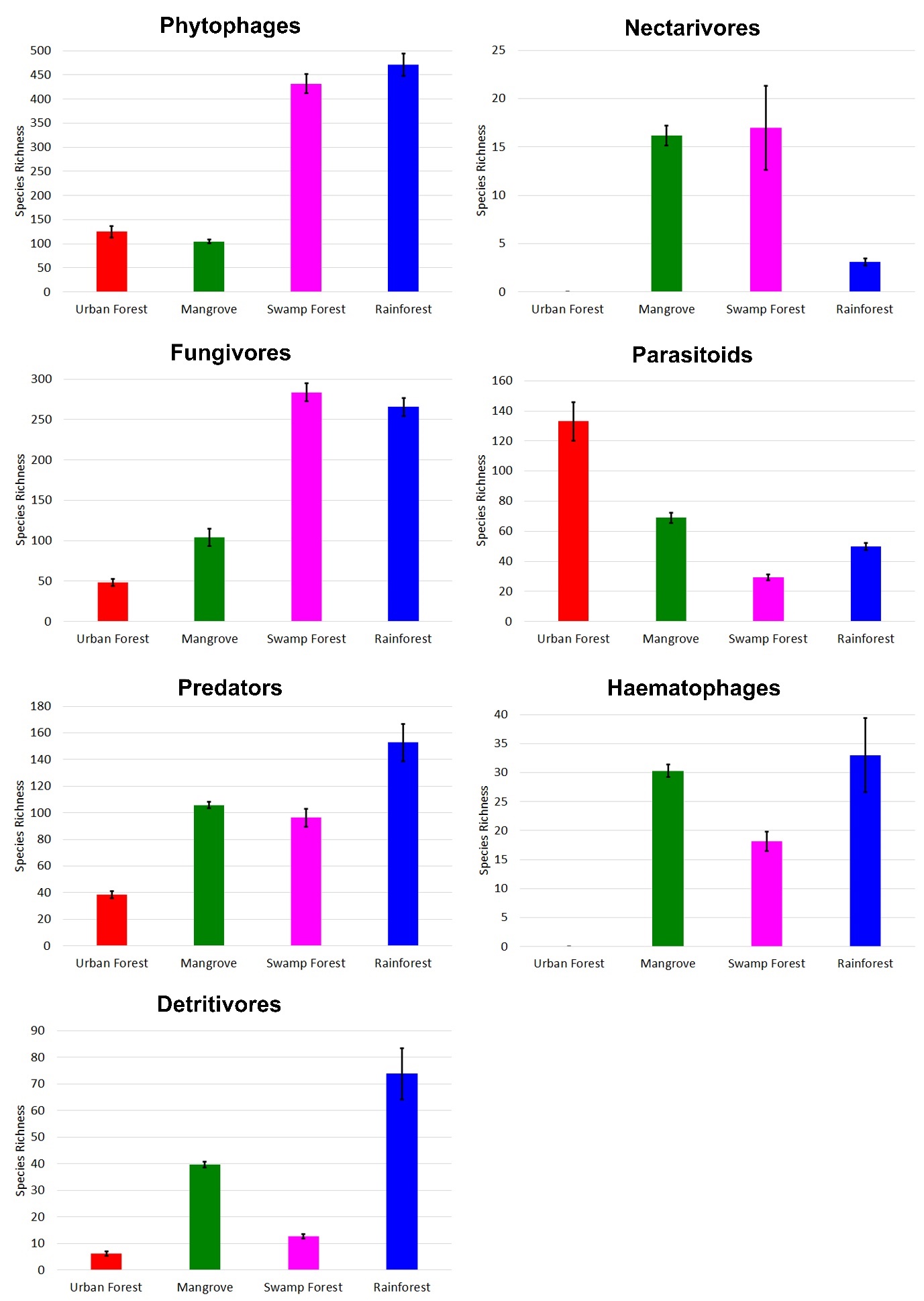

**Figure S7.** Comparison of species diversity across habitats (3% p-distance mOTUs) split by ecological guild. Values were taken at the point of rarefaction based on lowest coverage and include 95% confidence intervals. Mangrove sites are represented as a single habitat type.

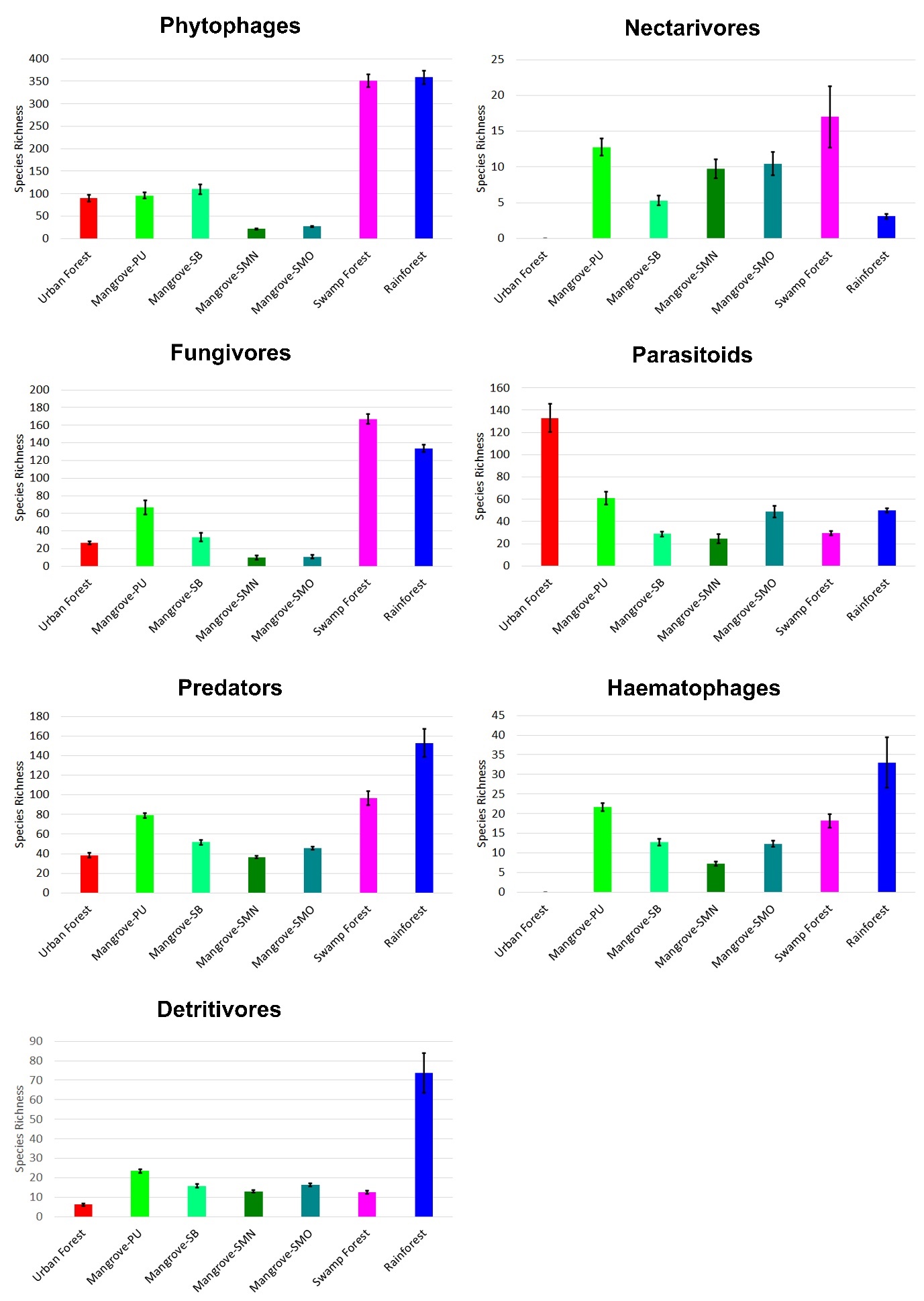

**Figure S8.** Comparison of species diversity across habitats (3% p-distance mOTUs) split by ecological guild. Values were taken at the point of rarefaction based on lowest coverage and include 95% confidence intervals. Mangrove sites are represented by Pulau Ubin (PU), Sungei Buloh (SB), Pulau Semakau old grove (SMO), Pulau Semakau new grove (SMN).

|  | No. of mOTUs |
| --- | --- |
| Dolichopodidae HK | 97 |
| Dolichopodidae SG | 88.2 |
| Dolichopodidae BR | 63 |
| Dolichopodidae HK + SG | 4.3 |
| Dolichopodidae HK + BR | 0 |
| Dolichopodidae SG + BR | 25.3 |
| Dolichopodidae All | 2 |
| Phoridae HK | 99 |
| Phoridae SG | 125.2 |
| Phoridae BR | 48 |
| Phoridae HK + SG | 11.9 |
| Phoridae HK + BR | 2 |
| Phoridae SG + BR | 11 |
| Phoridae All | 4.7 |
| Mycetophilidae HK | 23 |
| Mycetophilidae SG | 37.4 |
| Mycetophilidae HK + SG | 2 |

**Figure S9.** High species diversity and turnover for mangroves from Singapore, Brunei, and Hong Kong based on three Diptera families. Singapore data are rarefied to specimen numbers from Brunei and HK (error bars = 95% confidence intervals).
